# Lipid–ASO therapeutics exhibit differential tissue targeted delivery upon systemic or local CNS administration

**DOI:** 10.64898/2026.08.28.747711

**Authors:** Samantha Roudi, H. Yesid Estupiñán, Osama Saher, Cristiana Barradas, Emma Inganäs, Hoang-Ngoan Le, Nicolai Frengen, Radoslaw Grochowski, Svetlana Pavlova, Robert Månsson Welinder, Joel Z. Nordin, Annabelle Biscans, Pär Matsson, Rula Zain, Rickard Sandberg, Michael Hagemann-Jensen, Samir El Andaloussi

**Affiliations:** Department of Laboratory Medicine, Unit for Biomolecular and Cellular Medicine, Karolinska Institutet, Stockholm, Sweden; Karolinska ATMP center, Karolinska Institutet, Karolinska University Hospital, Stockholm, Sweden; Department of Cellular Therapy and Allogeneic Stem Cell Transplantation (CAST), Karolinska University Hospital, Stockholm, Sweden; Departamento de Ciencias Básicas, Universidad Industrial de Santander, Bucaramanga, Colombia; Department of Pharmaceutics and Industrial Pharmacy, Faculty of Pharmacy, Cairo University, Cairo, Egypt; Unit for Pharmacokinetics and Drug Metabolism, Department of Pharmacology, Institute of Neuroscience and Physiology, University of Gothenburg, Gothenburg, Sweden; Nucleic Acid Therapeutics, Discovery Sciences, BioPharmaceuticals R&D, AstraZeneca; Department of Laboratory Medicine, Division of Clinical Immunology, Karolinska Institutet, Stockholm, Sweden; National Genomics Infrastructure, Science for Life Laboratory, KTH Royal Institute of Technology, Stockholm, Sweden; Department of Clinical Immunology and Transfusion Medicine (KITM), Karolinska University Hospital, Stockholm, Sweden; SciLifeLab Gothenburg; Centre for Rare Diseases, Department of Clinical Genetics and Genomics, Karolinska University Hospital, Stockholm, Sweden; Department of Cell and Molecular Biology, Karolinska Institutet, Stockholm, Sweden

**Keywords:** Antisense oligonucleotides, lipid–ASO conjugates, drug delivery, systemic administration, intracerebroventricular administration, central nervous system

## Abstract

Antisense oligonucleotides (ASOs) are a powerful therapeutic modality, but their full potential is hindered by pharmacokinetic properties that affect tissue and cellular delivery. Lipid conjugation is increasingly used to modulate ASO’s biodistribution and promote extrahepatic activity, yet lipid-dependent effects on *in vivo* functional delivery, particularly in the central nervous system (CNS), remain less explored. Here, we performed a side-by-side *in vivo* comparison of cholesterol, palmitic acid (C16:0), docosanoic acid (C22:0), and eicosapentaenoic acid (C20:5) conjugated to a fully phosphorothioated 3-10-3 LNA gapmer ASO targeting the *Malat1* long non-coding RNA. Lipid–ASO conjugates were administered systemically or locally in the brain of mice and evaluated for tissue-level and cellular-level distribution by imaging, qPCR and single-cell RNA sequencing, simultaneously annotating cell origin and global transcriptional changes within the cell.

Following systemic administration in mice, lipid conjugation improved overall multi-organ efficacy compared to unconjugated ASO, but with pronounced tissue-specific differences. Single-cell sequencing of liver and heart transcriptomes revealed lipid-dependent cellular uptake patterns and transcriptional responses distinct from administration of unconjugated ASO. After intracerebroventricular administration, selected fatty acid conjugates enhanced silencing in deep brain regions such as the striatum, whereas cholesterol conjugation impaired functional delivery despite increased CNS retention. Light-sheet microscopy showed restricted parenchymal penetration of cholesterol-ASOs compared with broader but heterogeneous distribution of palmitic acid conjugate. Together, these findings demonstrate that lipid identity critically determines ASO efficacy, productive cellular uptake, and regional CNS engagement, emphasizing the need for context-specific lipid design in ASO therapeutic development.

**Highlights:**

- SC, fatty acid conjugates primarily potentiate hepatic over extrahepatic silencing
- SC, cholesterol conjugate confines silencing activity to the liver
- Single-cell sequencing reveals lipid conjugate-induced differential gene expression
- First study reporting evaluation of lipid-ASO functional activity in the CNS via ICV delivery
- Chol-ASO retention at CSF-facing membranes correlates with poor parenchymal activity

**Graphical abstract:** 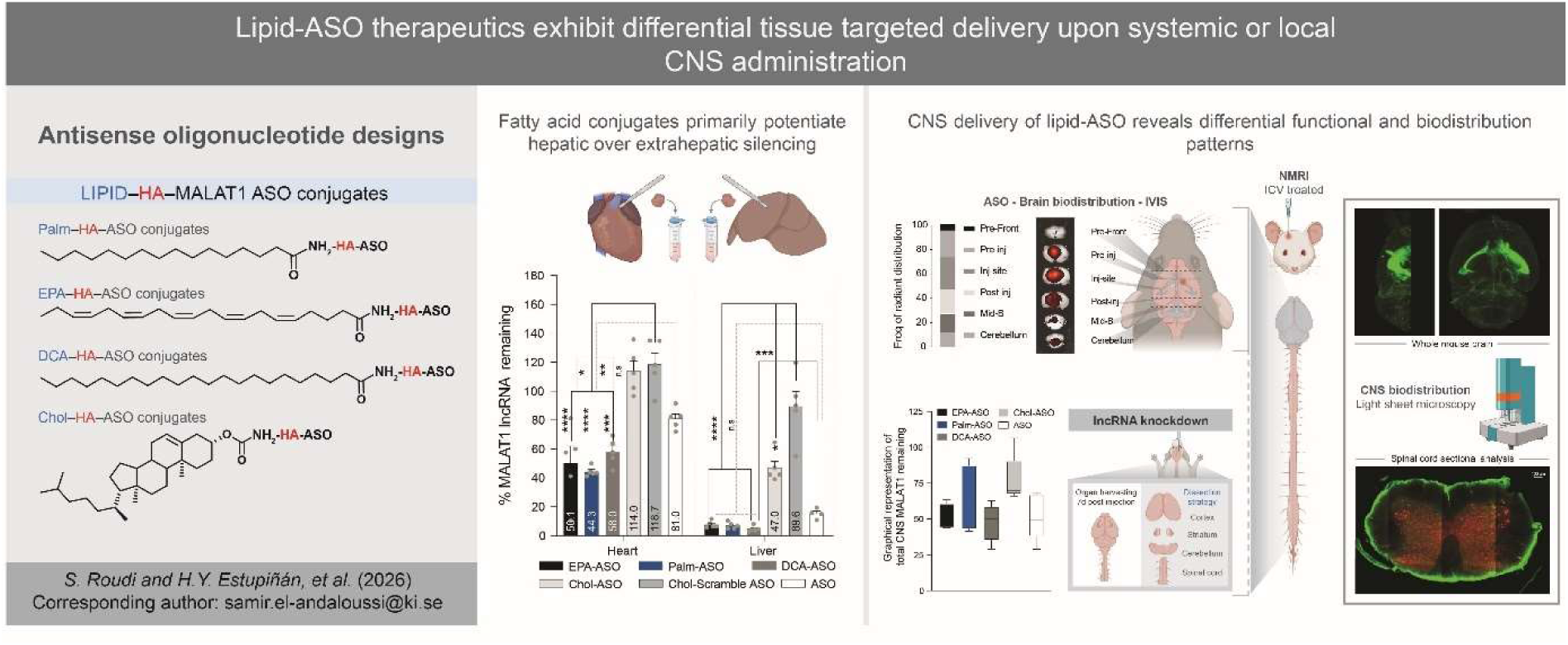

## 1. Introduction

Accelerated progress in the development of oligonucleotide (ON) therapeutics, such as antisense oligonucleotides (ASOs) and small interfering RNA (siRNA), can largely be attributed to their, in principle, straightforward yet highly specific drug design, which is dictated by the sequence of the disease-associated target RNA. However, their current status as disease-modifying drugs required overcoming several key challenges related to metabolic stability, pharmacokinetics, and pharmacodynamics. These limitations have been progressively addressed through multiple generations of chemical modifications to the oligonucleotide backbone, sugar, and nucleobase [1].

ASOs are short, chemically modified oligonucleotides, typically 15–21 nucleotides in length and composed of DNA- or RNA-like building blocks or hybrid chemistries within a single molecule [2]. Their therapeutic versatility arises from their adaptable chemical design, which enables multiple RNA-targeting mechanisms operating in different cellular compartments. In this study, ASOs refer specifically to gapmer chemistry, a design in which a central DNA gap flanked by chemically modified RNA-like wings recruits RNase H1 to induce degradation of target transcripts in nuclear or cytoplasmic compartments [3]. Gapmer ASOs frequently incorporate phosphorothioate (PS) backbone modifications, which confer increased nuclease resistance and increased affinity for plasma proteins, a property that reduces rapid renal clearance and promotes cellular uptake, thereby providing an intrinsic delivery advantage without the need for an additional targeting or delivery moiety [4].

The pharmacokinetic behavior of ASOs *in vivo* depends strongly on their chemical composition. Following systemic administration, ASOs rapidly and extensively bind to plasma proteins and distribute preferentially to organs with fenestrated or discontinuous capillary endothelium, resulting in predominant accumulation in the liver and kidneys. [5]. Consequently, most subcutaneously administered gapmer ASOs approved to date target hepatic diseases. In contrast, delivery to extrahepatic tissues such as skeletal muscle, heart, or the central nervous system (CNS) remains more challenging. Many neurological disorders would benefit from ASO-based therapies; however, ASOs do not readily cross the blood–brain barrier (BBB) after systemic administration [6,7]. To circumvent the BBB, several ASO drugs are delivered directly into the cerebrospinal fluid (CSF) via intrathecal (IT) injection. Nevertheless, heterogenous distribution, rapid drug clearance from the CSF and redistribution to peripheral tissues still limits local CNS persistence of the administered dose [8]. Collectively, these pharmacokinetic constraints highlight the need for strategies that improve ASO delivery to extrahepatic tissues following systemic administration or prolong retention and improve penetration at sites of local delivery.

A promising strategy to enhance ON therapeutic performance is conjugating them to targeting ligands, a well-established approach in siRNA delivery. The leading example is the triantennary *N*-acetylgalactosamine (GalNAc₃) ligand, which binds the asialoglycoprotein receptor (ASGPR) abundantly expressed on hepatocytes. Originally developed for siRNA therapeutics, GalNAc conjugation has subsequently been successfully translated to ASOs, with significantly enhanced hepatocyte uptake and potency. For example, the transthyretin-targeting GalNAc-ASO eplontersen achieves robust target silencing at a lower and less frequent dosing regimen than the unconjugated ASO inotersen, while maintaining favorable clinical outcomes with improved quality of life [9–12].

Building on the success of GalNAc-mediated liver targeting, a range of alternative ligand-conjugated ASO strategies are being explored to enable delivery to extrahepatic tissues. These include peptide-conjugated ASOs designed to enhance uptake in skeletal and cardiac muscle, pancreatic β-cells or the brain [13–19], antibody–oligonucleotide conjugates (AOCs) targeting receptors such as transferrin receptor 1 for muscle and brain targeting [20–22], and lipophilic ligands including cholesterol or fatty acids [23–26] that can increase tissue exposure and alter biodistribution profiles. By promoting receptor-mediated uptake or modulating interactions with circulating proteins and membranes, such ligand conjugates aim to overcome the intrinsic liver- and kidney-biased distribution of systemically administered ASOs and thereby improve therapeutic potency in otherwise difficult-to-reach tissues.

Here, we sought to improve the pharmacological performance of ASOs in extrahepatic tissues through lipid conjugation and to evaluate how different lipids influence ASO functional delivery and biodistribution. Four types of lipids were selected based on previous lipid–siRNA work and emerging reports on lipid-modified ASOs, demonstrating improved tissue uptake and altered pharmacokinetic profiles [23–27]. Using these conjugates, we evaluated how lipid attachment affects albumin binding, functional delivery, and tissue distribution of ASOs following both systemic administration and local delivery to the CNS.

Target knockdown was quantified at the RNA level across tissues to evaluate functional delivery. To characterize biodistribution following systemic administration, ASOs were labeled with a Cy3 fluorophore and visualized using In Vivo Imaging System (IVIS). To further investigate how lipid conjugation influences cell type–specific uptake patterns and gene regulation, we performed single-cell RNA sequencing on Cy3-positive cells isolated from liver and heart.

In addition to systemic delivery, we examined the performance of lipid-conjugated ASOs following direct administration into the cerebrospinal fluid. ASOs were administered intracerebroventricularly (ICV) and brain was dissected into distinct anatomical regions to evaluate functional target knockdown. Biodistribution within the CNS was first assessed using IVIS imaging and subsequently characterized at higher spatial resolution using light-sheet fluorescence microscopy, enabling three-dimensional visualization of ASO distribution across brain structures. Together, this work provides a side-by-side comparison of sterol, saturated and polyunsaturated fatty acids conjugated to ASOs across systemic and CNS delivery routes and offers insights into how lipid ligands influence tissue distribution, cellular uptake, and functional gene silencing.

## 2. Materials and methods

### 2.1 Antisense oligonucleotide conjugate synthesis

Reagents were purchased from Sigma-Aldrich part of the Merck group, TCI chemicals, WuXi AppTec, and used as received. Acetonitrile used for oligonucleotide synthesis was purchased as LiChrosolv grade from Sigma-Aldrich and further purified and dried by pushing through a MBRAUN SPS Compact using nitrogen gas immediately prior to use.

HA-*Malat1* (**10**), HA-Scramble, and *Malat1* ASO (**6**) were purchased from IDT and were used as received. Gapmer antisense oligonucleotides targeting human and mouse *Malat1* RNA used in experiments were ordered from IDT: fully phosphorothioated (PS) oligonucleotide backbone with following nucleotide sequence: <u>GCA</u>TTmCTAATAGmC<u>AGC</u>, where ten nucleotides in the middle are DNA, and are spanned by three locked nucleic acids (LNAs) on each end (underlined). mC denotes modification of cytosine base to 5-methylcytosine. For scramble control, the same chemistry was applied but with the following nucleotide sequence: <u>GGC</u>mCAATAmCGmCmCG<u>TCA</u>.

Cy3-labled sequences were synthesized on an ÄKTA OligoPilot Plus 10 synthesizer (GE Healthcare), on a 32 µmole scale, using a standard synthesis cycle of detritylation (3% dichloroacetic acid in toluene), coupling (coupling agent: 0.25 M 5-[3,5-Bis(trifluoromethyl)phenyl]-1H-tetrazole solution in acetonitrile), capping (Cap A: 20% N-methylimidazole and 80% acetonitrile; Cap B: 20% pyridine, 20% acetic anhydride and 60% acetonitrile), oxidation (0.05 M iodine in pyridine and water) or thiolation (0.2 M xanthane hydride in pyridine), and solid supports (UNY Primer Support 5G ∼353 μmol/g, GE Healthcare). All fully protected β-cyanoethyl phosphoramidite monomers were dissolved in anhydrous acetonitrile (0.1 M) under argon immediately prior to use. The phosphoramidite re-circulation time/coupling time for DNA monomers was 5 min and was extended to 10 min for LNA monomers. Stepwise coupling efficiencies and overall yields were determined by automated trityl cation absorption monitoring exceeding 98% for all oligonucleotides synthesized. At the end of the synthesis, the solid-support bound oligonucleotide was treated with ammonia (aq. 26%) solution (16 mL) at 35 °C for 18 h, and subsequently dried on a Speedvac. The crude mixture was purified by preparative HPLC (see general procedure).

Conjugates **1–5** and **8–9** were synthesized from the corresponding oligonucleotides bearing a 5′-hexylamino linker via reaction with the appropriate NHS ester, following general procedures. The crude mixtures were purified by HPLC and subsequently desalted to afford the desired products. Identity and purity of oligonucleotides was confirmed by liquid chromatography-mass spectrometry (LC-MS) using an Acquity I-class LC system, equipped with a PDA and coupled to an RDa via heated electrospray (BioAccord) by the Waters Corporation.

General procedure: A solution of oligonucleotide containing 5′-hexylamino liner (0.02 mmol) in water (2 mL) and triethylamine (0.048 mL, 0.35 mmol) at 60 °C was treated with a solution of lipid NHS-ester (2eq., 0.07 mmol)) in THF (1.000 mL) at 60 °C. The mixture was mixed at 60 °C OVN before all the volatiles were removed to afford the crude product. The crude mixture was purified by preparative HPLC using an XBridge C18 column (5 µm, 19 × 150 mm). The mobile phases consisted of 50 mM aqueous NH₄HCO₃ (solvent A) and methanol (solvent B). Elution was performed using a linear gradient of 20–100% solvent B.

### 2.2 Albumin binding assay

Kinetic experiments were performed at +25 °C with a WAVEDelta instrument (Malvern Panalytical) [28]. The running buffer was phosphate-buffered saline (PBS, pH 7.4). A PCH sensor chip was conditioned by injections of borate buffer (100 mM sodium borate, pH 9.0, containing 1 M NaCl; Xantec). Human serum albumin (HSA) was diluted in 10 mM sodium acetate buffer (pH 5.2) to 5 µg mL⁻¹. The sensor surface was activated by injecting a freshly prepared mixture of 1-ethyl-3-[3-dimethylaminopropyl]carbodiimide hydrochloride (EDC) and N-hydroxysuccinimide (NHS) (Xantec) for 420 s at 10 µL min⁻¹. HSA was then injected at 10 µL min⁻¹ into three of four channels until a surface density of approximately 4000 pg mm⁻² was reached, after which remaining reactive groups were blocked by injecting 1 M ethanolamine-HCl (Xantec) for 420 s. One channel served as a reference and was subjected only to the activation and deactivation steps (no HSA immobilization). The surface was allowed to equilibrate for at least 2h before binding experiments.

Surface functionality was assessed before and after the experiment by measuring binding of the established ligand diclofenac to immobilized HSA, using multi-cycle kinetic mode with a two-fold dilution series starting at 20 µM. Literature reports two diclofenac binding sites on HSA, with KD values in the range 2-20 µM [29]. Equilibrium analysis of the sensorgrams using a 1:1 binding model yielded apparent KD values in the range 6–31 µM.

Antisense oligonucleotides were diluted in PBS to 5–10 µM. Kinetic measurements were performed using the RAPID (Repeated Analyte Pulses of Increasing Duration) [30] scheme with the “Weak Binders” setting (400 µL min⁻¹; 5 s association; 20 s dissociation; 40 Hz data acquisition). After each sample injection, the surface was regenerated with 25% (v/v) ethylene glycol (80 µL min⁻¹, 30 s injection followed by 120 s dissociation). Buffer blanks (PBS, pH 7.4) were injected before each sample. A solution of 0.5% (v/v) DMSO in PBS (pH 7.4) was injected every tenth sample and used for solvent (DMSO) correction.

Sensorgrams were solvent-corrected and blank-subtracted in WAVEcontrol (version 4.9). Data were fitted using the heterogeneous binding model with the traditional kinetics engine.

### 2.3 Animal studies

All the animal experiments were approved by The Swedish Board of Agriculture (Jordbruksverket) and carried out in compliance with their guidelines. Male and female NMRI mice were used in all experiments unless otherwise stated. Mice were bought from Janvier labs and housed at the Preclinical Laboratory (PKL), Novum, Karolinska University Hospital, Huddinge, under specific pathogen-free conditions. Mice were allowed to acclimatize for one week before the start of the experiment. Husbandry and housing conditions were compliant with national animal welfare legislation and were monitored daily by animal care staff.

### 2.4 Systemic administration of lipid-conjugates

For subcutaneous (SC) administration, NMRI mice (N=5 mice/ group) were administered with the compound at the dose 50 mg/kg and sacrificed 48h post injection, followed by organ collection (heart, lung, liver, kidneys, spleen) and snap freezing of organs at -80°C until further processing.

For intravenous (IV) administration, NMRI mice (N=4-5 mice/group) were administered with the Cy3-conjugated compounds at the dose 10 mg/kg into tail vein. Mice were sacrificed 48h post injection, followed by organ collection (brain, heart, lung, liver, kidneys, spleen) and ex-vivo IVIS imaging of Cy3 to assess biodistribution.

For single cell sequencing experiment, NMRI mice (N=3/group, and N=2 for Vehicle control group) were administered IV Cy3-conjugated compounds at the dose 5 mg/kg into tail vein. Mice were sacrificed 48h post injection, heart and liver were dissected, washed in PBS and stored in PBS+1% FBS on ice until further processing.

### 2.5 Intracerebroventricular administration of lipid-conjugates

For intracerebroventricular (ICV) delivery of lipid-ASO conjugates, mice were anesthetized with isoflurane and positioned on a stereotaxic frame with prewarmed heating pad to maintain body temperature at 37°C. The eyes of mice were protected with eye ointment and mice were injected SC with buprenorphine (0,1 mg/kg) prior the procedure. Next, the head was shaved and a small incision made to expose bregma to measure the coordinates for injection into right lateral ventricle (anteroposterior 0.3 mm, mediolateral 1 mm, dorsoventral 3 mm). Mice were injected with 5 µL of 10 μg ASO conjugate diluted in 0,2 μm filtered PBS at the injection rate 1 μL/min and the Hamilton Neuros syringe was left inserted for another 2 min before slow withdrawal. The bone was closed with the bone wax and the incision sutured.

For functional delivery experiment, mice (N=4/group) were sacrificed 7 days post-injection. Spinal cord and brains were collected, and brains were dissected into four anatomically distinct regions – cortex, striatum, cerebellum and rest, which encompass of midbrain, thalamus, hypothalamus, hippocampus and other minor regions. Brain regions were frozen on dry ice and stored at -80°C until further processing.

For biodistribution studies, mice were administered with 5 µL of 10 μg Cy3-labeled conjugates were administered and sacrificed 48h post-injection for ex vivo IVIS (N=3/group) and 3D imaging (N=1/group).

### 2.6 RNA isolation

For systemically administered mice, total RNA from organs was isolated with TRI Reagent® (T9424, Sigma-Aldrich). First, 1 mL of TRI Reagent® was added to the 2mL tubes containing tissue and 5 mm steel bead, and homogenized using Qiagen TissueLyser II, followed by centrifugation at 12,000 x g at 4°C for 5 min. Chloroform was added to the clear supernatant in 1:5 v/v ratio and samples were vigorously shaken for 1 min before centrifugation at 12,000 x g for 10 min at 4°C for phase separation. The aqueous upper phase was collected, followed by the addition of an equal amount of isopropanol, mixing by inversion and incubation at -20°C overnight. The precipitated RNA was centrifuged at 4°C, 12,000 x g for 10 min, and the RNA pellets washed two times with 70% ethanol before solubilization in RNAse free water and incubation at 55°C for 10 min. Total RNA was measured on NanoDrop and stored at -80°C before cDNA synthesis.

For ICV administered mice, total RNA for collected brain regions was isolated using Maxwell® RSC simplyRNA Tissue Kit (AS1340) according to manufacturer’s instructions. Briefly, brain regions were homogenized in 200 µL of chilled 1-Thioglycerol/Homogenization Solution using Qiagen TissueLyser II and the homogenate was loaded into cartridges in assigned wells. DNase I solution, RSC plungers and elution tubes with Nuclease-Free Water were added/positioned in the cartridge/deck trays as per instruction and simplyRNA Tissue method was selected on Maxwell instrument for RNA isolation. Total RNA was measured on NanoDrop and stored at -80°C before cDNA synthesis.

### 2.7 qPCR quantification of *Malat1* long non-coding RNA

cDNA synthesis was performed using Applied Biosystems™ High-Capacity cDNA Reverse Transcription Kit (43-688-13). RT-qPCR reactions for *Malat1* detection were prepared using TaqMan™ Fast Advanced Master Mix for qPCR as per manufacturer’s instructions. Mouse *Malat1* was detected using TaqMan assay (Assay ID: Mm01227912_s1, FAM dye) and data normalized to housekeeping gene 36B4/RPLP0 with forward primer sequence GAGGAATCAGATGAGGATATGGGA, reverse primer sequence AAGCAGGCTGACTTGGTTGC and probe sequence TCGGTCTCTTCGACTAATCCCGCCAA (JOE dye). RT-qPCR detection was performed on StepOnePlus Real-Time PCR System with following settings: 50°C for 2 min, polymerase activation at 95°C for 2 min, PCR cycle at 95°C for 3 sec and 60°C for 30 sec, repeated 40x. Relative gene expression of *Malat1* was determined by ΔΔCt method and normalized to the vehicle-treated control group.

### 2.8 Ex vivo IVIS imaging

For systemic biodistribution, mice were injected IV with Cy3-labeled lipid-ASO conjugates at a dose 10 mg/kg, sacrificed 48h post-injection and organs (brain, heart, lung, liver, kidneys, spleen) were collected for ex-vivo IVIS imaging immediately after harvesting. An IVIS® Spectrum imaging system (PerkinElmer, Waltham, MA, USA) was used for ex vivo fluorescence imaging with a 570/620 nm excitation/emission filter pair. Total radiant efficiency ([p/s]/[µW/cm²]) was quantified within a defined region of interest (ROI). All images were acquired and analyzed using Living Image® 4.4 software (PerkinElmer, Waltham, MA, USA).

For CNS biodistribution, mice were injected ICV with 5 µL of 10 μg Cy3-labeled lipid-ASO conjugates as described above. 48h post-injection mice were anesthetized with isoflurane, followed by terminal heart puncture. Chest was open to access the heart for transcardial perfusion with 25 mL of PBS+ 0,002% heparin, followed by 25 mL of 4% PFA. Whole brain and spinal cord were further fixed in 4% PFA at room temperature (RT) overnight. Brain was dissected into following segments: pre-frontal, pre-injection, injection-site, post-injection, mid-brain and cerebellum. A representative segment from cervical and lumbar regions of the spinal cord were also harvested. Ex vivo IVIS imaging was performed on PFA-fixed brain regions, with instrument setting as described above.

### 2.9 Whole brain and spinal cord clearing

Mice were sacrificed 48h post-injection and perfused as described above. Whole central nervous system dissection was performed as previously described [31], brain and spinal cord were further isolated and fixed in 4% PFA overnight at RT under gentle agitation. Next, CNS tissues were cleared using Miltenyi Biotech’s MACS Clearing Kit (Cat Nr.: 130-136-719) and following corresponding protocol entitled “Immunostaining and clearing of mouse brain hemispheres with preservation of endogenous fluorescent protein signal”[32]. Briefly, whole brain and spinal cords were incubated with permeabilization solution for 24h at RT, followed by dehydration with increasing percentage of tert-butanol, starting from 30% to final 99,5% tert-butanol at 28°C. Next, samples were cleared with Clearing Solution provided in MACS Clearing kit at RT overnight and replaced with fresh Clearing Solution for additional 6h. Samples were stored in MACS Imaging Solution (Cat Nr.: 130-126-335) and 1% triethylamine at RT until imaging. The protocol was modified only for spinal cord samples, where an extra step including immunostaining and washing was added. VioR667 NeuN (Cat Nr: 130-131-153, clone REA1131) and VioR667 TH (Cat Nr: 130-131-157, clone REA1159) were used at 1:50 dilution.

### 2.10 Light sheet microscopy

Lightsheet microscopy and 3D image acquisition was performed in ethyl cinnamate (ECi) using a light-sheet fluorescence microscope (Ultramicroscope Blaze, Miltenyi Biotec, Germany) equipped with 1.1x (NA. 0.1), 4x (NA 0.35), and 12x (NA.0.53) objective lenses. The system provides an axial resolution of approximately 0.5-5 μm.

Cy3 signals were detected using 561nm laser diode and emission filter 620/60 nm filter and for spinal cords, VioR667 signals for NeuN and TH were detected using 639nm laser for excitation and 680/30 emission filter on PCO Edge 4.2 MP camera.

Image acquisition was performed with ImSpector software (v7.1) followed by automatic data processing by MACSiQ view – 3D large volume (v1.1) software. Cy3 signals were detected by excitation with 561nm laser diode and emission filter 620nm /60 filter and PCO Edge 4.2 MP camera. We selected the resolution we needed which was delivered by the overview 1x, 4x or 12x objective and 1x zoom.

### 2.11 Preparation of single cell suspensions from liver and heart

Single cells were prepared using Miltenyi Biotech’s dissociation kits and gentleMACS Octo Dissociator with Heaters (130-095-937). Heart was dissociated following the protocol “Dissociation of adult mouse heart using the Multi Tissue Dissociation Kit 2 (Cat. No. 130-110-203), and liver following the protocol “Liver Dissociation Kit, mouse (Cat. No. 130-105-807). As for deviation from the protocol, cell suspensions of both tissues were filtered through 100 μm cell strainer (Falcon™ 352360). Additionally, red blood cell lysis was performed for liver, following the same protocol as in Multi Tissue Dissociation Kit 2 but without PBS + enzyme A washing step. All samples were stored in PBS+1%FBS on ice until sorting and further diluted in PBS containing DAPI (Thermo Scientific™, Cat. No. 62248, diluted to 1 μg/mL) just before the cell sorting.

### 2.12 Single cell sorting

Cells were sorted on BD Discover S8 spectral image-based sorter at MedH Flow Cytometry Core Facility that receives funding from the Infrastructure Board at Karolinska Institutet. To apply the gating on healthy cell populations, DAPI was used to discern live/dead cells in combination of bright-field imaging of the cell sorter. Vehicle treated samples were sorted from DAPI negative population, while additional Cy3 gating was applied for Cy3-label containing samples. Single cells were sorted through a 100 µm nozzle into 384-well Armadillo PCR plate (Thermo Scientific) containing 0,3 µl Smart-seq3xpress lysis buffer [33]. Plates were sealed with aluminum Axygen Microplate Sealing Film, spun down immediately and stored at -80 °C until further processing.

### 2.13 Smart-seq3xpress library generation and sequencing

Sorted single cells were prepared for Smart-seq3xpress library preparation as previously described [33]. In brief, sorted single cells from heart and liver were reversed transcribed and amplified using 14 cycles. 1uL of the diluted amplified cDNA was subsequently tagmented using 0.005uL TDE1 (Illumina) per well, before index PCR with custom Illumina Nextera compatibable index primers for 14 cycles. Afterwards libraries were pooled, bead-cleaned and evaluated via Qubit (Thermo Scientific) and Bioanalyzer (Agilent). Single-cell RNA sequencing libraries were sequenced on MGI DNBSEQ G400 platform using universally compatible (App-D) sequencing reagents according to the manufacturer’s instructions.

### 2.14 Single cell data preprocessing, read alignment and transcript quantification

Raw FASTQ files were processed using zUMIs (v2.9.7)23. Two batches of data were processed separately. In the YAML configuration both batches were run with find pattern: ATTGCGCAATG and base definition: UMI(12–21); cDNA(25–100). Batch 1 read 2 was parsed with cDNA (1–100) and barcode (BC, 101–120). UMIs were collapsed using a Hamming distance of 1, and barcode binning was set to 1. Reads were aligned to the mouse reference genome (mm39 using STAR24(v2.7.3a) with gene annotations from GENCODE GRCm39 (Release 29). Additional STAR parameters included --clip3pAdapterSeq CTGTCTCTTATACACATCT. Gene-level read and UMI count tables were generated, capturing counts from exons and introns. The two batches were combined post preprocessing.

### 2.15 Single-cell RNA-sequencing data processing, annotation, and lipid-ASO specific analysis

Raw count matrices were imported into R QC’ed and processed using Seurat. Cells of low quality were excluded based on dataset-specific quality control thresholds (mapped to exon+intron > 30% of reads, contain more than 10.000 sequenced reads, and contain less than 20% mitochondrial reads). Genes detected in less than 10 cells were removed prior to downstream analyses. Where relevant, metadata including sample identity, tissue, condition, mouse replicate, and fluorescence-based uptake measurements (MFI) were incorporated into the Seurat object. Data were normalized and scaled using standard Seurat workflows. Highly variable genes (3000 features) were identified and used for dimensionality reduction. Principal component analysis (PCA) was performed, followed by nearest-neighbor graph construction, unsupervised clustering, and visualization using Uniform Manifold Approximation and Projection (UMAP), following Seurat default parameters. Cell types were assigned manually based on established marker genes and the top differentially expressed genes for each cluster. Annotation was first performed at a broad lineage level, and labels were added back to the Seurat object and used for all downstream cell type resolved analyses.

Cell type composition was quantified per sample and compared across conditions to assess whether lipid-ASO treatment altered the relative abundance of major populations. To distinguish transcriptional state changes from compositional changes, differential expression analyses were performed within annotated cell types by comparing treatment conditions to control cells from the same tissue and cell type. Analyses were restricted to cell types with sufficient cell numbers > 20 cells per cell type per condition to support robust testing. Single-cell differential expression testing was performed using MAST with sequencing depth as latent variable. Uptake-associated effects were examined by incorporating matched fluorescence intensity measurements as per-cell metadata and comparing their distributions across tissues and cell types. Expression of *Malat1* was analyzed across conditions and annotated cell populations to assess target-associated responses.

### 2.16 Statistical analysis

Statistical analysis was performed using ordinary oneway ANOVA (Alpha: 0.05) followed by Dunnett’s multiple comparisons test to compare treatment groups against the vehicle control for each tissue separately.

## 3. Results and discussion

### Identification of lipids for *Malat1* ASO conjugation, synthesis and protein binding properties

To evaluate the impact of lipid conjugation on ASO biodistribution and activity *in vivo*, we conjugated selected lipids to a model 3-10-3 LNA-DNA-LNA gapmer ASO with fully phosphorothioated (PS) backbone targeting the long non-coding RNA metastasis associated lung adenocarcinoma transcript 1 (*Malat1*), a well-established preclinical target owing to its robust and relatively uniform expression across tissues [13,14]. Lipids were conjugated via NHS ester chemistry to ASO bearing a 5′-hexylamino linker [9,34]. The selection of lipids for ASO conjugates included cholesterol as a representative for steroids, palmitic acid (Palm; C16:0) and docosanoic Acid (DCA; C22:0) as representatives for long saturated fatty acids, and eicosapentaenoic acid (EPA; C20:5) as a representative for polyunsaturated fatty acid (**Figure 1A and Supplementary Table 1)**. Building on previous studies of lipid-conjugated oligonucleotides, we investigated their effects across multiple systemic organs rather than focusing on a single target tissue, enabling direct comparison of tissue distribution and functional delivery in the liver, heart, lungs, kidneys, and spleen.

**Figure 1.**
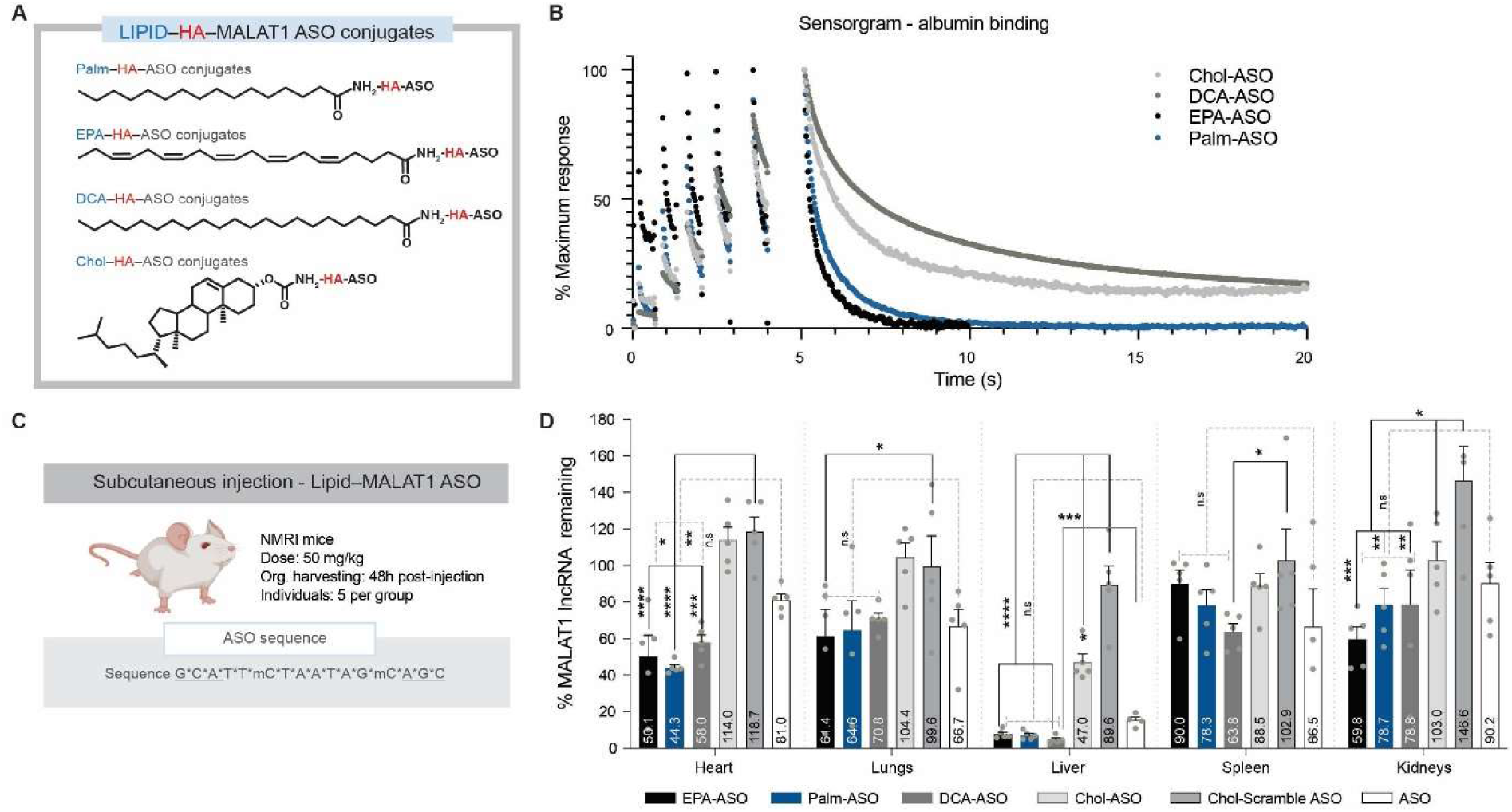
Lipid–ASO conjugates and functional *in vivo* delivery upon subcutaneous administration. **A)** Schematic representation of the selected lipid-ASO conjugates, where lipid components are conjugated to the 5’-end of a *Malat1* targeting ASO via hexylamine (HA) linker. Palm = palmitic acid (C16:0); DCA = docosanoic acid (C22:0); EPA = eicosapentaenoic acid (C20:5); Chol = cholesterol. **B)** Kinetics of lipid-ASO binding to albumin (HSA), measured using grafting-coupled interferometry. Sensorgrams were normalized to each compound’s maximum response to facilitate comparison of the dissociation rate profiles. **C)** Experimental details of systemically administered ASOs and *Malat1* ASO sequence **D)** *Malat1* knockdown in indicated organs 48h after single subcutaneous injection of 50 mg/kg lipid-ASO or ASO only in female NMRI mice (N=5). *Malat1* transcripts were quantified by RT-qPCR, normalized to a housekeeping gene *RPLP0*, and the data presented as percentage of remaining *Malat1* transcript in vehicle control. Each dot represents an individual animal; bar height and error bars indicate the group mean ± standard deviation (SD). Significance for multiple variables comparison was calculated using ordinary oneway ANOVA, heart (F_5,24_ = 22.60, ****p =<0.0001); liver (F_4,19_ = 57.32, ***p =<0.0001) and Kidney (F_5,24_ = 5.151, **p =0.0024).

When administered systemically, the biodistribution of PS-modified ASOs is dictated by their association with multiple plasma proteins. Lipid conjugation alters the plasma binding profiles and consequently, the ASO’s biodistribution and pharmacological activity. We therefore first assessed the impact of lipid conjugation on binding to the most abundant plasma protein, serum albumin (SA), using a grating-coupled interferometric biosensor assay analyzed with a two-site binding model. Sensorgrams revealed slower dissociation kinetics for DCA and cholesterol conjugates than for Palm and EPA, which both dissociated rapidly and completely from albumin (**Figure 1B)**. The dissociation kinetics were reflected in the calculated dissociation constants (KD), with all lipid conjugates displaying measurable albumin affinity, in contrast to the unconjugated ASO for which no binding was detected. Affinity for the higher-affinity binding site ranked as follows: DCA> Chol> Palm > EPA, with mean residence time (MRT; calculated as the reciprocal of the dissociation rate constant) following the same order (**Supplementary table 2**). Taken together, these findings demonstrate that lipid identity critically determines the strength and persistence of albumin engagement, consistent with reports that conjugation of hydrophobic moieties to ASOs shifts their protein binding profile towards albumin and lipoproteins, with downstream consequences for tissue distribution and cellular uptake [24,25,35].

### RNA knockdown in organs after subcutaneous administration of lipid-ASO conjugates

We next investigated *Malat1* lncRNA downregulation in NMRI mice after a single SC-administered dose (50 mg/kg) of lipid-ASO conjugates and its non-conjugated counterpart (**Figure 1C**). The injected dose was well tolerated, and mice showed no apparent signs of acute toxicity. Heart, lungs, liver, spleen and kidneys were collected 48 h post-ASO administration and knockdown was assessed using RT-qPCR. Collectively, all *Malat1* targeting ASOs induced the most potent RNA silencing in liver. Chol-ASO showed liver-specific targeting, with no activity in other organs, except for the spleen with modest 11.5% target silencing. While cholesterol conjugation served as a reference for hepatic delivery and achieved slightly more than 50% target KD, surprisingly, free ASO as well as all other lipid-ASO conjugates resulted in significantly (p≤0.0001) greater target suppression in liver: approx. 85% with free ASO and between 92-95% with EPA, Palm and DCA conjugates, respectively (**Figure 1D**). This is consistent with a study by Østergaard *et al*., who directly compared cholesterol- and palmitate-conjugated ASOs administered IV and SC; IV delivery of both conjugates produced strong hepatic and cardiac muscle activity exceeding unconjugated ASO, whereas SC administered Chol-ASO showed reduced muscle activity, pointing to route-dependent differences in tissue exposure and suggesting that cholesterol conjugates may be less efficiently mobilised from the SC depot into systemic circulation [34].

While our results demonstrated enhanced ASO potency in the liver upon lipid conjugation, our primary aim was to compare different lipids side-by-side for their functional delivery to extrahepatic organs following SC administration. In the heart, palmitic acid showed significantly (p≤0.0038) enhanced potency compared to ASO only, consistent with previous reports showing its strong cardiac and skeletal muscle delivery properties [5]. Additionally, in heart, EPA- and DCA-ASO still exhibited potent target silencing compared to unconjugated ASO, although the difference was significant (p≤0.0159) only for the EPA-ASO conjugate. In lungs, Palm-, EPA- and DCA-ASOs performed similarly to unconjugated ASO, resulting in around 35% target downregulation. This same degree of activity was observed in the spleen for unconjugated and DCA-conjugated ASO, while other lipid conjugates were not as efficient in target silencing. As reported previously, kidneys are the primary clearance organ and thus accumulate ASOs but the degree of functional delivery is not correlated to the degree of accumulation [5] – this poor bioactivity for most conjugates as well as unconjugated ASO was also observed in our experiment (**Figure 1D** and **2A**). While unconjugated ASO silenced around 10% of *Malat1* in kidneys, DCA and Palm induced around 20% silencing. EPA exhibited the highest functional delivery efficiency among all lipids, achieving more than 40% target knockdown, which is consistent with the bioactivity profile previously reported for siRNA conjugates [27,36].

**Figure 2:**
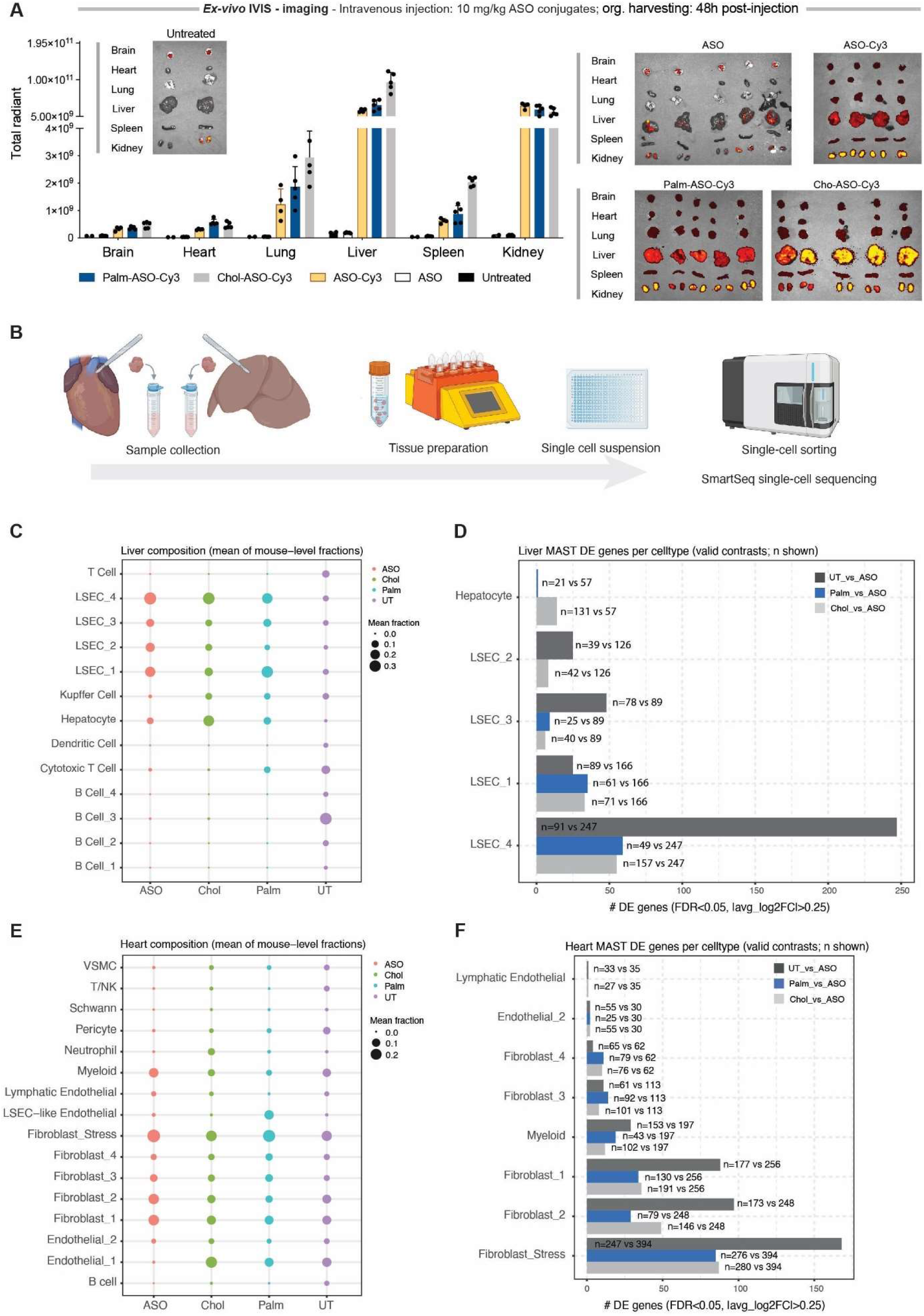
Uptake of selected lipid-ASOs as determined in bulk tissue and on a single-cell level. **A)** Ex vivo IVIS imaging following systemic administration of Cy3-labeled ASOs (10 mg/kg). Left – total radiant efficiency of Cy3 signal quantified by defined region of interest (ROI = whole organ). Each dot represents an individual animal; bar height and error bars indicate the group mean ± standard deviation (SD). Right – Cy3 signal in individual organs captured by IVIS. **B)** Schematic representation of experimental workflow for single cell sequencing of liver and heart following systemic administration of Cy3-labeled ASOs (5 mg/kg). **C)** Cell-type composition of liver samples **D)** Number of differentially expressed genes on a single-cell level in liver samples, analyzed using MAST. ASO group was compared to other groups. **E)** Cell-type composition of heart samples **F)** Number of differentially expressed genes on a single-cell level in heart samples, analyzed using MAST. ASO group was compared to other groups.

As shown in the results above, the capacity of lipid–ASOs to mediate functional delivery varies across organs, indicating that ligand design should be tailored to the specific disease context. Although systemically administered ASOs are predominantly active in the liver, many diseases would benefit from improved systemic distribution and activity. Therefore, we aimed to determine whether lipid conjugation could enhance the overall systemic efficacy of ASOs. To estimate the cumulative activity of each conjugate, we summed the mean percentages of Malat1 lncRNA remaining for all organs and calculated the average, thereby obtaining a measure of conjugate efficacy independent of organ-specific differences. As expected, the cholesterol conjugate displayed the lowest overall ASO activity, consistent with its strong liver specificity. In contrast, Palm, EPA, and DCA conjugates showed approximately 10% greater target knockdown on average compared to the unconjugated ASO, corresponding to ∼48% versus ∼57% *Malat1* remaining, respectively (**Supplementary Fig. 1**). These findings suggest that conjugation with selected lipids can improve the systemic performance of ASOs following systemic administration, which is particularly relevant for diseases requiring broad, multiorgan engagement.

### Biodistribution poorly reflective of functional delivery

Having characterized the differences in functional delivery of lipid–ASO conjugates, we next investigated whether these could be attributed to variations in biodistribution. Chol- and Palm-ASOs were selected as representative conjugates with contrasting pharmacodynamic profiles and compared to the unconjugated ASO. To avoid confounding effects of SC administration, particularly local depot formation and absorption kinetics, we administered Cy3-labeled ASOs IV, ensuring immediate systemic exposure and allowing organ accumulation to more directly reflect intrinsic tissue affinity. The IV dose was reduced five-fold relative to the SC dose (10 mg/kg) to achieve comparable peak plasma concentrations, consistent with the SC bioavailability of approximately 54–78% reported for mipomersen, a well-characterized PS-ASO [37]. Animals were sacrificed 48 hours post-injection, after which organs were analyzed by *ex vivo* IVIS fluorescence imaging.

Consistent with the established biodistribution of PS-modified ASOs, the primary sites of ASO accumulation were the liver and kidneys, with substantially lower signals detected in the lungs, spleen, heart, and brain. Among the conjugates, Chol-ASO showed higher accumulation in liver, lungs, and spleen relative to Palm-ASO and unconjugated ASO. Notably, this contrasts with the comparatively modest and hepatically confined functional activity of Chol-ASO observed upon SC dosing and pointing once more to administration route-dependent differences in systemic exposure. Following SC injection, cholesterol conjugates have been shown to accumulate extensively at the injection site, which limits their systemic bioavailability. This has been documented for both Chol-ASO and Chol-siRNA conjugates [34,38,39]. Kidney accumulation was high across all groups, as expected; however, the disconnect between renal signal intensity and functional activity is well-established and reflects the predominance of non-productive, lysosome-directed uptake in proximal tubule cells rather than pharmacologically active intracellular delivery [5].

### Single-cell sequencing reveals distinct uptake profiles and cellular responses in liver and heart

The aforementioned results underscore the limitations of using bulk tissue accumulation as a predictor of pharmacological activity and highlight the importance of distinguishing between productive and non-productive uptake at a cell-type-specific level. To further investigate the impact of lipid conjugation on cellular distribution, we performed single-cell RNA sequencing following IV administration of Cy3-labeled ASOs of liver and heart, as organs with the highest silencing activity.

Preliminary experiments indicated saturation of Cy3 signal at an IV dose of 10 mg/kg (data not shown). Therefore, a reduced dose of 5 mg/kg was selected to better resolve differences in cellular uptake associated with lipid conjugation. Chol-ASO-Cy3, Palm-ASO-Cy3 and unconjugated ASO-Cy3 were administered and single cell suspensions from liver and heart were prepared 48h post injection using a commercial dissociation protocol. Cells from treated animals were sorted based on Cy3 fluorescence to isolate ASO-positive cells (**Figure 2B**). Since untreated mice do not contain Cy3-positive cells, control samples were processed using the same dissociation protocol and sorted based only on DAPI exclusion to establish the baseline cell-type composition recovered by the dissociation procedure. The clustering of the single cells divided by cell types and treatment groups is presented by UMAP in **Supplementary Figure 2A.**

In untreated liver samples, the recovered cell population consisted predominantly of non-parenchymal cells, with immune populations, including Kupffer cells, accounting for approximately 80% of recovered cells, while the remaining cells were largely liver sinusoidal endothelial cells (LSECs) with only a small fraction of hepatocytes (**Figure 2C**). This distribution is consistent with the expected enrichment of non-parenchymal cells resulting from the dissociation protocol used, which is designed to preferentially recover these populations and may underrepresent hepatocytes.

In contrast, the cell-type composition of the Cy3-positive fraction from ASO-treated mice differed markedly from this baseline. Among ASO-positive cells, LSECs and hepatocytes together accounted for more than 80% of the sorted population (and over 95% in the lipid-unconjugated ASO group), whereas immune cells represented a much smaller fraction. Because the dissociation protocol was identical for all samples, this shift reflects preferential association or uptake of Cy3-labeled ASOs by specific cell populations rather than differences in cell recovery. Within the Cy3-positive population, the proportion of hepatocytes was modestly higher in the Chol-ASO-Cy3 group (∼30%) than in the Palm-ASO-Cy3 and unconjugated ASO-Cy3 groups (∼20%). Given the low baseline representation of hepatocytes following dissociation, this relative enrichment suggests that cholesterol conjugation increases hepatocyte-associated uptake, consistent with previous reports using cholesterol-conjugated ASOs containing a hexylamine succinimide linker [40]. Notably, Kupffer cells accounted for a measurable fraction of Cy3-positive cells in both Chol- and Palm–ASO-Cy3 groups (∼10%), whereas they were nearly absent in the unconjugated ASO-Cy3 group (<0.01%) (**Figure 2C)**. Together, these results indicate that the majority of ASO-positive cells are liver sinusoidal endothelial cells, with comparatively smaller differences observed in hepatocyte and Kupffer cell fractions. However, this observation should be interpreted with caution, as it likely reflects differences in uptake within the recovered cell population rather than absolute cell type– specific uptake. To date, relatively few studies have investigated the cellular distribution of ASOs in vivo, and these have primarily relied on histological approaches, including immunofluorescence, immunohistochemistry, or more recently in situ hybridization combined with image analysis, as well as comparisons between isolated hepatocyte and non-parenchymal cell fractions. Single-cell transcriptomic approaches integrated with direct measurements of ASO uptake remain rare, and therefore the extent to which different methodologies capture the true in vivo cellular distribution of ASOs has yet to be fully established [41–44].

Next, we sought to assess productive versus non-productive uptake at the single-cell level by integrating transcriptomic data with corresponding fluorescence intensity measurements recorded during cell sorting. However, variability in *Malat1* transcript levels was high even in untreated samples, and treated samples did not exhibit sufficient knockdown to enable a reliable correlation between ASO uptake (as measured by Cy3 signal intensity) and target suppression (**Supplementary Fig. 2B)**. One possible explanation for the limited silencing observed is the relatively low dose used in this experiment, which may result in insufficient intracellular ASO concentrations and hence target engagement. In addition, the early time point may further limit detectable activity, as only a small fraction of internalized ASO is expected to escape endosomal compartments and reach its site of action.

Although no substantial *Malat1* knockdown was observed, we further hypothesized that the exposure duration and extent of cellular uptake might still be sufficient to induce measurable transcriptomic changes. To assess cell type–specific responses to lipid-conjugated ASOs, differential expression analyses were performed comparing ASO-positive cells treated with Chol-ASO or Palm-ASO to those treated with unconjugated ASO. In addition, comparisons to untreated controls were included. Single-cell differential expression testing was conducted using MAST, and analyses were restricted to cell types with sufficient representation (n>20 cells). In the liver, the LSEC_4 population exhibited the highest number of differentially expressed genes (DEGs); however, the overall response remained modest (**Figure 2D**). Compared to unconjugated ASO, both Chol-ASO and Palm-ASO treatments resulted in approximately 55 DEGs. In contrast, comparison to untreated samples revealed a larger transcriptional response for unconjugated ASO (∼240 DEGs). Across other cell types, the number of DEGs was highly cell type–dependent but generally low to moderate, ranging from 10 to 50 genes across groups of comparisons.

The same analysis was performed on heart samples (**Figure 2E**). In contrast to the liver, cell type composition was not markedly altered between untreated and treated groups. As expected based on the dissociation protocol, samples were enriched for non-parenchymal cells, predominantly fibroblast subtypes and endothelial cells. Notably, the Endothelial_1 population was absent among ASO-positive cells in the unconjugated ASO group, despite representing a substantial fraction of cells in all other groups, including untreated samples (∼10–20%). This observation suggests that unconjugated ASO is not efficiently taken up by this endothelial subtype. As in the liver, we were unable to reliably distinguish productive from non-productive uptake at the single-cell level (**Supplementary Fig. 2B**). Differential expression analysis revealed the strongest transcriptional response in stressed fibroblasts, with approximately 80 differentially expressed genes (DEGs) following treatment with Chol-ASO or Palm-ASO compared to unconjugated ASO. In contrast, unconjugated ASO treatment elicited a somewhat stronger, though still modest, response (∼120 DEGs) when compared to untreated controls (**Figure 2F**). This trend, although with lower responses was also observed in additional fibroblast subtypes and myeloid cells.

Together, this data shows for the first time that the lipid conjugation itself induces transcriptional changes to the recipient cells that are different from the one elicited by naked ASO.

### Unilateral bolus ICV administration of lipid**–**ASO conjugates results in potent *Malat1*

**knockdown in deep brain regions and spinal cord.**

Next, we investigated the impact of lipid conjugation on CNS ASO delivery via Intracerebroventricular (ICV) administration, which, to our knowledge, has not been previously explored for lipid-conjugated ASOs. While intrathecal (IT) delivery is widely used clinically for CNS-targeted ASO therapies, emerging evidence suggests that distribution throughout deeper brain regions can be heterogeneous and limited [12,45]. We administered NMRI mice a single dose of lipid–ASO (10 ug, ∼0.5 mg/kg) directly into CSF via the right lateral ventricle. Mice were sacrificed 7 days post injection, after which spinal cord and brains were collected. Brains were dissected into four anatomically distinct regions: cortex, striatum, cerebellum and rest, the latter encompassing of midbrain, thalamus, hypothalamus, hippocampus and other minor regions (**Figure 3A**). Expression of *Malat1* was determined for both hemispheres, irrespective of injection site.

**Figure 3.**
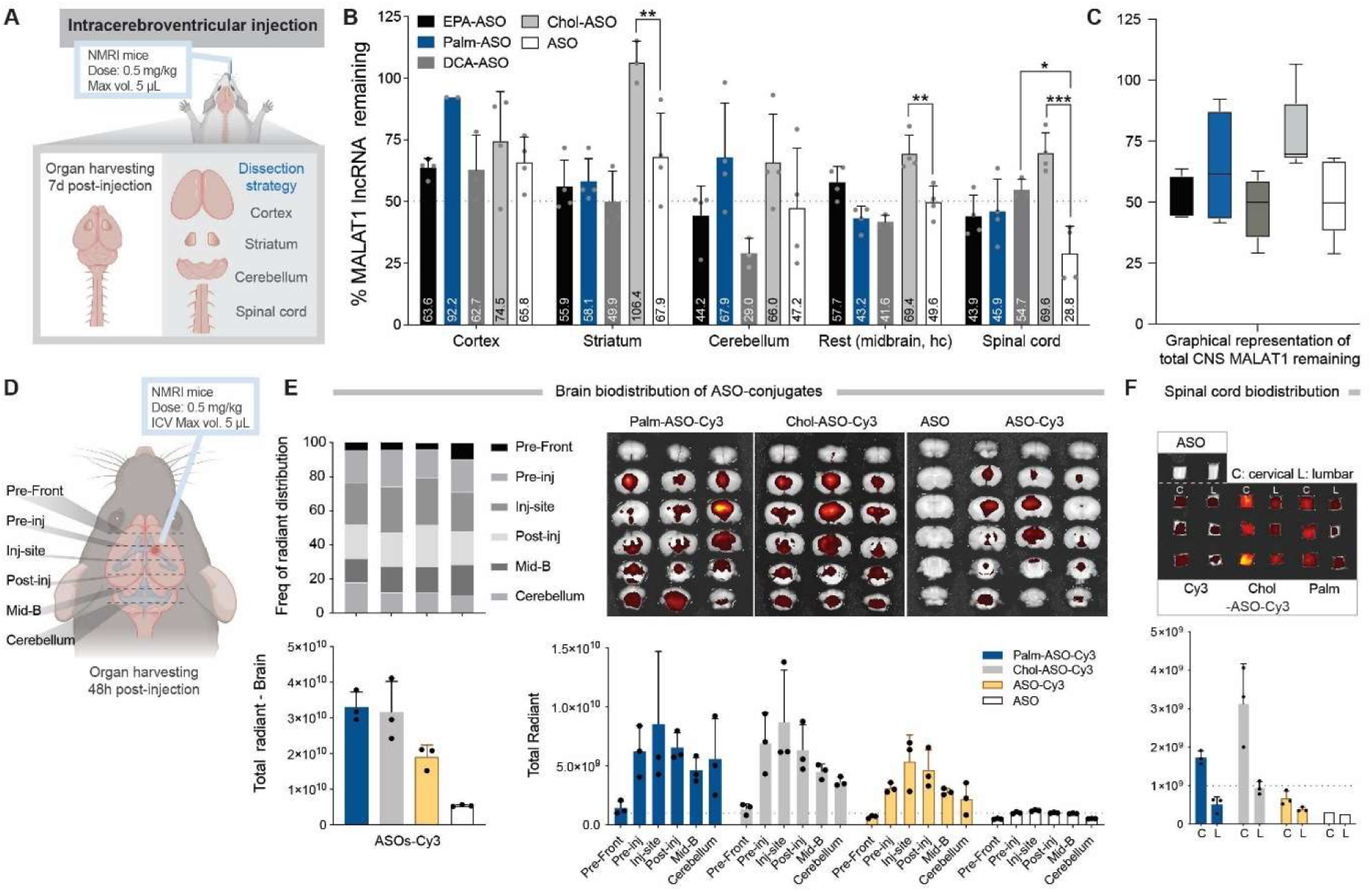
Functional delivery and biodistribution of lipid-ASO conjugates to the CNS. **A)** Schematic representation of experimental setup and brain dissection strategy after unilateral bolus ICV administration of 0.5 mg/kg lipid-ASO or ASO only in female NMRI mice. **B)** Percentage *Malat1* lncRNA remaining in CNS regions. Each dot represents an individual animal (N=3 or 4); bar height and error bars indicate the group mean ± standard deviation (SD). Significance was calculated using ordinary one-way ANOVA, striatum (F_4,13_ = 10.12, **p =0.0050); rest (F_4,14_ = 13.59, **p =<0.0014) and SC (F_4,14_ = 9.185, ***p =<0.0002). **C)** Graphical representation of cumulative activity of each conjugate, calculated by averaging the percentage of Malat1 lnRNA remaining for all CNS regions. **D)** Schematic representation of dissection strategy for ex vivo IVIS imaging of brains for selected lipid-ASO-Cy3 conjugates. **E)** Ex vivo IVIS images of coronal brain sections displayed from pre-frontal cortex to the cerebellum, section thickness = 1 mm. Bottom left - quantified total fluorescence signal captured across all six sections. Bottom right - signal intensity of individual sections. Each dot represents one animal, and the bar height represents the mean value of the group. **F)** Ex vivo IVIS images of cervical and lumbar sections of spinal cord and corresponding quantification of the fluorescence signal in the graph below.

ICV administration has been shown to achieve more effective ASO distribution to deep brain regions than intrathecal (IT) delivery in rodents [8,46–48]. We therefore used the ICV route to evaluate whether lipid conjugation influences ASO penetration and functional delivery to the striatum. Here, conjugation with DCA-, EPA- and Palm-ASO proved to be beneficial as they all showed improved target silencing potency compared to unconjugated ASO (from ∼50-56% vs ∼68% Malat1 lncRNA remaining, respectively). In cerebellum, similar patterns of improved activity were observed with DCA- and EPA-ASO, whereas in the cortex, the responses of the same compounds were comparable to those of the unconjugated ASO control. However, Palm conjugation did not appear to be beneficial for the delivery to cerebellum or cortex. In the “rest” tissue regions, the three conjugates performed varyingly, with Palm and DCA being more potent, and EPA less potent compared to unconjugated ASO. On the other hand, cholesterol conjugation resulted in suppression of functional delivery in all CNS regions we evaluated (**Figure 3B**). Activity in brain regions other than striatum appeared to be lipid conjugate dependent.

In the spinal cord, unconjugated ASO was markedly more active than the lipid conjugates, suggesting that lipid conjugation may either limit the movement of ASOs through the CSF to distal spinal regions or reduce their productive uptake by spinal cord cells once they arrive. Collectively, these data suggest that functional delivery of ASOs upon ICV administration is intrinsically lipid-dependent: while some lipid conjugates improve activity in deep brain regions such as the striatum, they often do so at the expense of potency in other injection-distant regions.

### Lipid-ASOs exhibit enhanced retention in deep brain

Given the variable performance of Palm-ASO in different regions and relatively scarce activity of Chol-ASO throughout the CNS, we next assessed their biodistribution to determine whether the results on ASO activity could be explained by different retention patterns of lipid-ASOs in the CNS. Mice were administered with a single ICV dose of ASO-Cy3, Chol-ASO-Cy3 or Palm-ASO-Cy3, or with non-labelled ASO to establish the imaging baseline, and were sacrificed 48h post injection. Whole brain and spinal cord were fixed overnight in 4% paraformaldehyde, followed by dissection as represented in **Figure 3D**. Sections were imaged with In Vivo Imaging System (IVIS) detecting Cy3 signal.

*Ex vivo* IVIS imaging and the corresponding quantitative analysis of total radiant efficiency in the brain demonstrate that Cy3 signal remained detectable across the CNS at 48 h post-injection. The signal was substantially higher for Chol-ASO-Cy3 and Palm-ASO-Cy3 in comparison to unconjugated ASO-Cy3, indicating enhanced retention of lipid-ASO conjugates at 48h. Across all three ASOs, the highest fluorescence signal was observed at the site of injection, with gradual decline towards both, rostral and caudal brain regions (**Figure 3E**, total radiant signal). A similar signal intensity pattern was observed in the spinal cord, where Chol-ASO-Cy3 yielded the highest Cy3 signal, followed by Palm-ASO-Cy3 and unconjugated ASO-Cy3. Expectedly, the lumbar spinal cord, the most injection-distant CNS region, exhibited lower signal intensities for all three ASOs relative to the cervical region (**Figure 3F**).

### 3D whole brain imaging reveals distinct biodistribution patterns of lipid-ASOs

While section-based IVIS measurements clearly demonstrate differences in overall signal intensity and rostrocaudal distribution, the method’s limited spatial resolution and the section thickness prevent accurate assessment of how the ASOs are distributed within the intact 3D CNS architecture. To overcome these limitations, we next employed light-sheet fluorescent microscopy (LSFM) on optically cleared whole brains and spinal cord to resolve the full three-dimensional distribution of the ASO. The experimental setup was the same as for *ex vivo* IVIS imaging experiment described above and representative slides are depicted in **Figure 4**.

**Figure 4.**
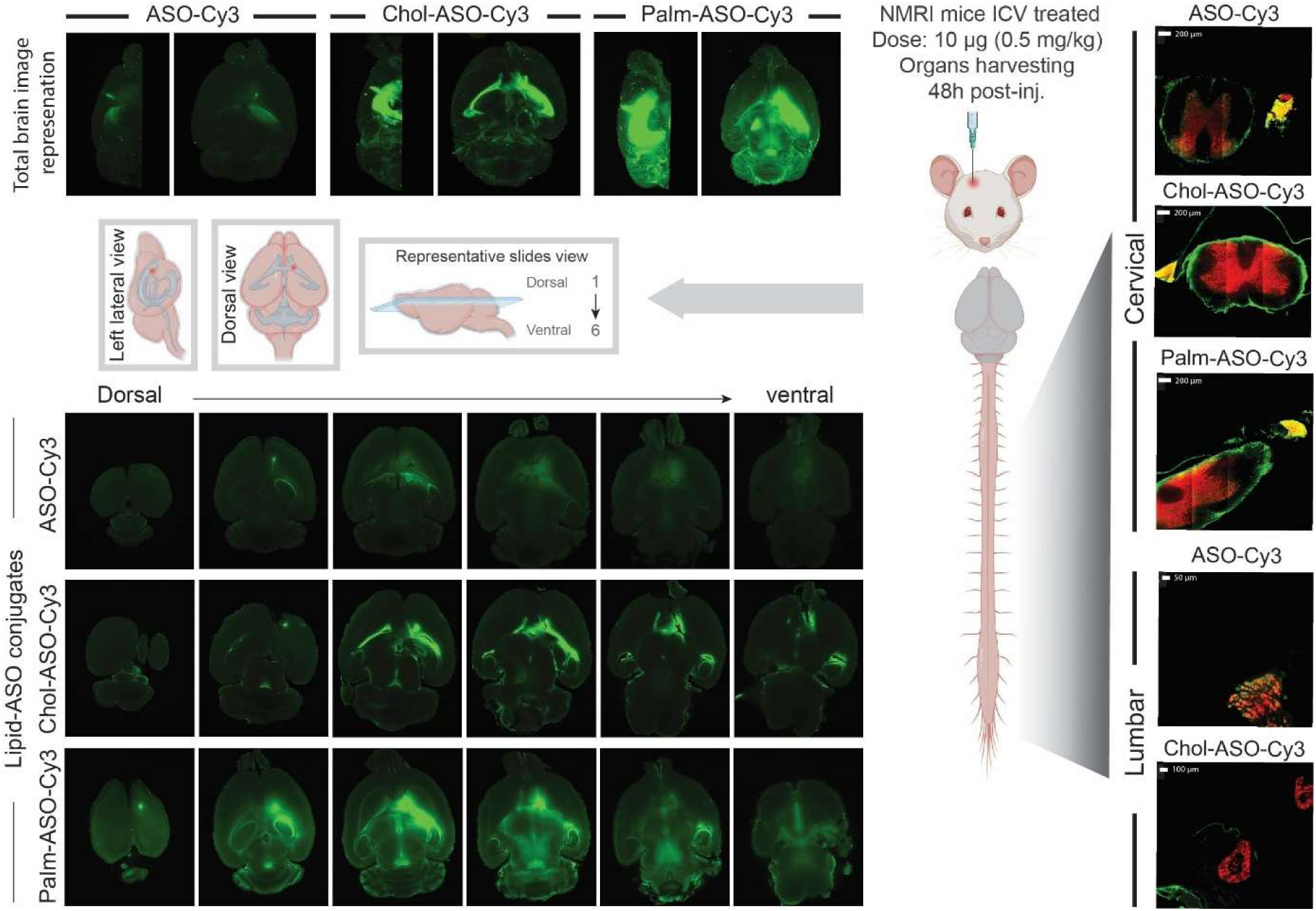
Biodistribution of lipid-ASO-Cy3 in the whole mouse brain and part of spinal cord upon ICV administration using light sheet microscopy. (Left and Right panels) Whole brain and spinal cord were isolated for clearing using the MACS® Clearing Kit with preservation of endogenous fluorescent signal. Image acquisition was performed using Ultramicroscope Blaze (Miltenyi Biotec, Germany). Cy3-labeled lipid-ASO conjugates (green), and additionally for spinal cords anti-NeuN-VioR667and anti-TH-VioR667 (red). Snapshots of whole brain and spinal cord are represented here.

At 48 hours, all three molecules (Cy3-labeled unconjugated ASO, Chol-ASO and Palm-ASO) remained detectable, although the fluorescent signal from the unconjugated ASO-Cy3 was only faint, consistent with a more rapid clearance from the CNS compared to the lipid-conjugated variants, as was also observed by *ex vivo* IVIS imaging. Using LSFM, we were able to resolve distinct distribution patterns between the Chol-and Palm-conjugated ASOs. **Figure 4A** shows lateral and dorsal projections of total fluorescence, where Chol-ASO-Cy3 displays highly localized Cy3 intensity with strong contrast relative to the background, indicating restricted diffusion into surrounding parenchyma. Spatially, the signal is confined yet robust across both hemispheres and closely follows the morphology of the lateral ventricular system. This pattern suggests efficient bilateral ventricular diffusion but limited penetration beyond ventricular surfaces. Such restricted parenchymal access is consistent with the poor functional activity in distal regions observed for Chol-ASO and aligns with previous observations reported for cholesterol-conjugated siRNAs [49].

In contrast, Palm-ASO-Cy3 exhibited similarly strong fluorescence intensity but reduced contrast relative to background, presenting a more diffuse distribution than Chol-ASO-Cy3. The spatial pattern remained strongly influenced by the injection site into the right lateral ventricle, while still displaying bilateral dissemination into regions corresponding to the “rest” (midbrain, hippocampus, thalamus, etc.) and cerebellum, matching the areas dissected for functional analyses. Based on this biodistribution profile and its strong signal in the right hemisphere, knockdown observed in the striatum and cortex for Palm-ASO in **Figure 3A** is likely driven primarily by proximity to the injection site, whereas effects in the “rest” and cerebellum reflect distribution into both hemispheres.

### Poor parenchymal penetration of lipid-ASOs in the spinal cord

To evaluate ASO biodistribution along the neuraxis following ICV administration, the spinal cord was harvested together with the whole brain at 48h post ICV administration of 10 ug Cy3-labeled ASO. Spinal cords were additionally immunostained for two established neuronal markers present in CNS: Neuronal Nuclei (NeuN), a nuclear and perinuclear marker of post-mitotic neurons, and Tyrosine Hydroxylase (TH), a cytoplasmic marker of dopaminergic neurons (**Figure 5, Supplementary Video 1**).

**Figure 5.**
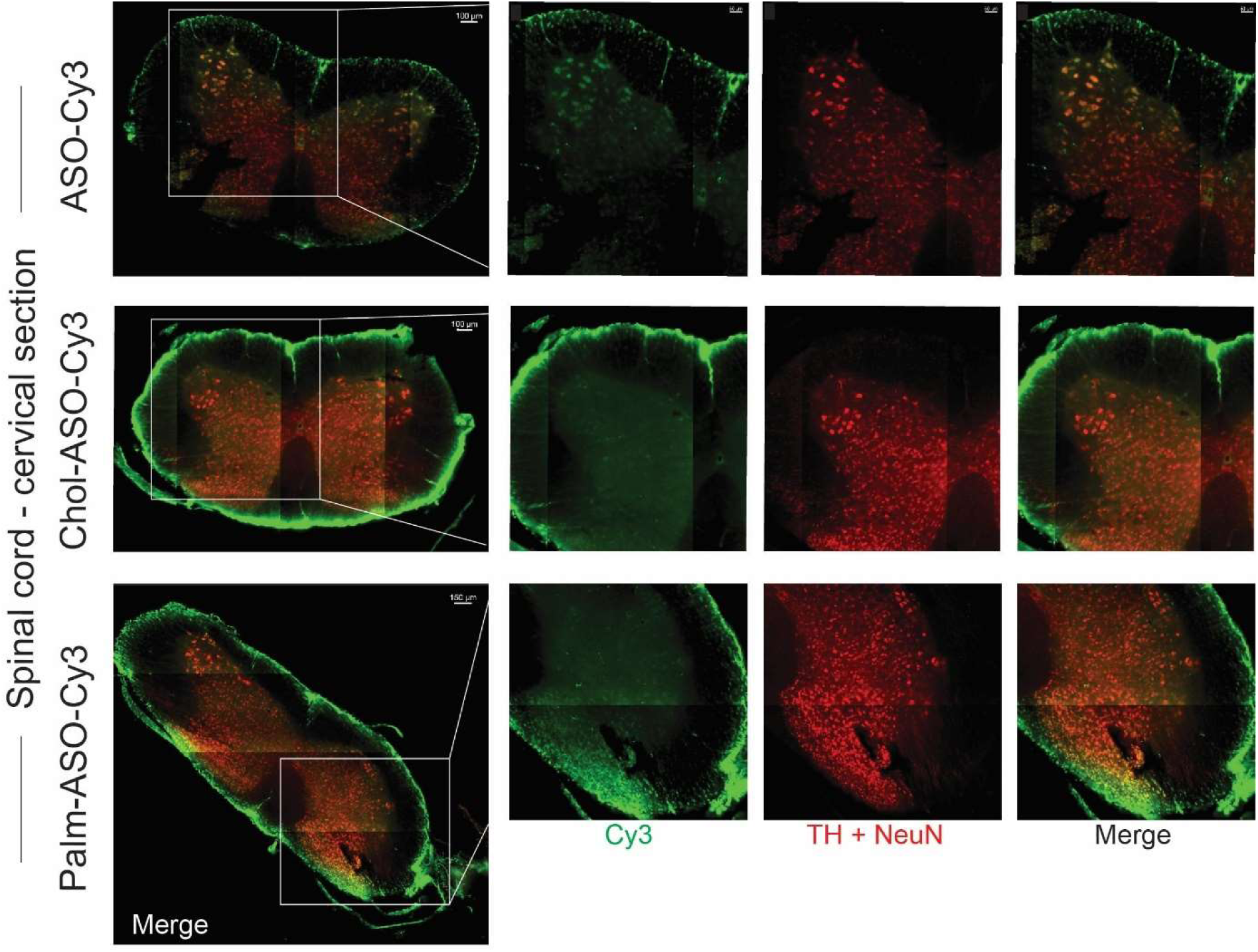
Biodistribution of lipid-ASO-Cy3 in cervical spinal cord upon ICV administration using light sheet microscopy. Spinal cords were harvested 48h post ICV administration and optically cleared using the MACS® Clearing Kit with preservation of endogenous fluorescent signal with additional immunostaining for NeuN and TH markers. Cy3-labeled lipid-ASO conjugates (green), and anti-NeuN-VioR667and anti-TH-VioR667 (red). Close up on the region of interest to display penetration beyond meningeal surfaces.

The overall Cy3 signal intensity in the spinal cord was lower for the unconjugated ASO than for the two lipid conjugated variants, indicating its lower retention in cervical spinal cord. Spatially, fluorescence signal wasstrongest at the outer surface of the spinal cord, consistent with the anatomical location of the pia mater, which forms the interface between the cerebrospinal fluid (CSF) and the spinal cord parenchyma. This pattern is in line with prior studies demonstrating rapid association of intrathecally or ICV-administered ASOs with meningeal surfaces and perivascular structures before parenchymal entry [50,51].

Despite this surface enrichment, ASO signal was also detectable within both white and gray matter regions of the spinal cord, where partial colocalization with NeuN- and TH-positive cells was observed, indicating intracellular neuronal association. This finding is consistent with reports showing neuronal and glial uptake of ASOs within 24–48 h following central administration [47,50,52].

In the case of Chol-ASO-Cy3, a markedly different distribution pattern was observed. Cholesterol-conjugated ASOs exhibited strong accumulation at the pia mater with minimal penetration into the spinal cord parenchyma, similar to the poor distribution seen in the brain. Signal within the white matter was sparse, and penetration into the gray matter was largely absent. This pronounced surface localization suggests adsorption to CSF-facing membranes and meningeal structures, which likely limits access to neuronal targets within the spinal cord. We therefore hypothesize that such retention at meningeal interfaces is a primary contributor to the reduced functional efficacy of Chol-ASOs across the CNS, including spinal cord.

Palm-ASO-Cy3 showed an intermediate distribution profile. While overall fluorescence intensity was slightly lower than that observed for the cholesterol conjugate, signal remained strongest at the pia mater but exhibited more apparent projections into the white matter. Penetration into the gray matter was largely restricted to the dorsal horn of the spinal cord and showed partial colocalization with neuronal markers. This pattern suggests that palmitic acid conjugation permits limited parenchymal entry relative to cholesterol conjugation, although penetration remains spatially constrained.

### Intracerebroventricularly administered lipid-ASOs distribute and are retained in the peripheral nervous system

Interestingly, for all three compounds, the strongest ASO signal within the cervical region of the neuraxis was observed in the peripheral nervous system, specifically within the dorsal root ganglia (DRG). Due to the high signal intensity, single-cell resolution could not be reliably achieved; however, ASO-positive cells appeared to co-localize with neuronal markers. While NeuN and TH are classically associated with CNS neurons, NeuN expression has also been reported in subsets of DRG neurons [53], supporting the validity of these observations.

In the lumbar region, co-localization of ASO signal with neuronal markers within the DRG was retained only in the unconjugated ASO-Cy3 group (**Supplementary Video 2A**). In contrast, DRG-associated signal was largely absent in the Chol-ASO-Cy3 group (**Figure 4 (right), Supplementary Video 2A**), suggesting reduced accessibility of cholesterol-conjugated ASO to lumbar DRG neurons. Unfortunately, DRG tissue from the Palm-ASO-Cy3 group was inadvertently lost during sample preparation and could not be evaluated.

Lumbar DRG represent a promising therapeutic target for ASO-based approaches in pain management due to their lack of a blood–brain barrier and high accessibility via intrathecal delivery. Numerous studies have demonstrated robust ASO-mediated gene suppression and analgesic efficacy in DRG following intrathecal or systemic administration [53–55]. Our observations suggest that lipid conjugation, and particularly cholesterol modification, may be suboptimal for achieving effective ASO distribution along the spinal neuraxis, especially in the lumbar spinal cord and DRG, which are frequent targets in pain-related indications.

Importantly, the dose used in this study lies at the lower end of those typically associated with functional target suppression in the CNS. Previous studies have demonstrated dose-dependent increases in parenchymal penetration and neuronal association of ASOs following central administration. Therefore, we cannot exclude the possibility that higher doses of lipid-conjugated ASOs may overcome meningeal retention and achieve improved penetration into spinal cord parenchyma and DRG neurons [47,50].

### Low protein binding capacity in CSF as a possible mechanism of high retention of lipid-ASOs at CSF-brain interface

While lipid conjugates have been explored to improve the extrahepatic bioavailability of oligonucleotide therapeutics through increased plasma protein binding, this exact mechanism might be the cause of the (partially) opposite effect observed in the context of local CNS delivery, owing to fundamental differences in protein abundance and composition between plasma and CSF. In mouse plasma, palmitoyl-conjugated ASOs display high affinity for albumin, while cholesterol-conjugated ASOs associate primarily with HDL and LDL, with substantial albumin binding also reported [23,25], interactions that collectively underpin their favorable systemic biodistribution. CSF, however, has around 150-fold lower total protein concentration compared to plasma (0.2–0.7 mg/mL vs. 60–70 mg/mL, respectively) [56], and given the approximate total CSF volume of 35-40 µL in adult mice [57], the absolute amount of carrier protein available is extremely limited. To provide a theoretical and simplified upper-bound estimate of carrier availability relative to drug load, we calculated the molar binding capacity of the two principal lipid-binding proteins in CSF, i.e. albumin and ApoE-containing HDL-like lipoprotein particles, at the experimental ICV dose of 10 µg (**Table 1**). Albumin fatty acid-binding site estimates were derived from human serum albumin crystal structures, which are considered broadly applicable across mammalian species given the conserved three-domain architecture of albumin, though mouse-specific binding site data are not available [58,59]. Both Palm-ASO-Cy3 and Chol-ASO-Cy3 were assumed to occupy up to seven hydrophobic binding sites per albumin molecule, consistent with the known affinity of both long-chain fatty acids and cholesterol moieties for the same hydrophobic pockets of albumin [23–25]. ApoE-lipoprotein intercalation capacity was estimated based on general HDL biophysical parameters, with mouse CSF ApoE concentration extrapolated from the microdialysis measurements in mice (∼6 µg/mL) [60]; however, no studies have directly measured HDL-like lipoprotein particle number or concentration in mouse CSF. Critically, when albumin binding is accounted for as the dominant carrier for both conjugates, the total carrier capacity is exceeded approximately 1.9-fold by both Palm-ASO-Cy3 and Chol-ASO-Cy3 at the 10 µg ICV dose, with ApoE-containing HDL-like lipoprotein contributing negligibly to total capacity relative to albumin.

**Table 1.** Theoretical estimation of carrier protein capacity in CSF. (a) ApoE copy number per CSF HDL-like particle has not been directly measured; estimate is extrapolated from plasma HDL ApoA-I stoichiometry (2–5 molecules per particle by MALDI-MS) as a structural analogy, given that ApoE is the primary scaffold protein of CSF HDL-like particles. (b) Free (unesterified) surface cholesterol accessible for exogenous intercalation is estimated at 5–10% of total HDL cholesterol content (∼46 molecules/particle on average); the remainder is esterified in the hydrophobic core and not accessible. This is a theoretical estimate; to our knowledge, no direct measurement of exogenous cholesterol intercalation capacity per CSF HDL particle exists in the literature.

|  | Parameter |  | Palm-ASO-Cy3 | Chol-ASO-Cy3 | Reference |
| --- | --- | --- | --- | --- | --- |
| Albumin | CSF volume | 35-40 μL | | | [57] |
|  | Concentration | ~0.195 mg/mL |  |  | [56] |
|  | Molecular weight [g/mol] | 66500 |  |  |  |
|  | Total (molar) | ~117 pmol |  |  | Calculated |
|  | Fatty acid binding sites | 7 |  |  | [58,59] |
|  | **Total carrier capacity** | 819 pmol | 819 pmol | 819 pmol | Calculated |
| ApoE-lipoprotein | Concentration (ApoE) | ~6 μg/mL | | | [60] |
|  | Molecular weight [g/mol] | 34200 |  |  |  |
|  | ApoE molecules per CSF HDL particle (a) | ~2-5 (a) |  |  | [61,62] |
|  | Lipoprotein particles (at 5 ApoE/particle) | ~1.4 pmol |  |  |  |
|  | Free cholesterol surface sites/particle (b) | ~3–5 accessible surface sites (~5–10% of ~46 total cholesterol molecules) |  |  | [63–65] |
|  | **Total carrier capacity (molar)** | ~4-7 pmol | Not accounted | 4-7 pmol | Calculated |
|  | **Total combined capacity** |  | ~819 pmol | ~823–826 pmol | Calculated |
|  | ASO MW (g/mol) |  | 6287.59 | 6461.83 | This study, Supp Table 2 |
|  | **ASO molar load at 10 μg ICV dose** |  | ~1,590 pmol | ~1,548 pmol | Calculated |
|  | **Drug:carrier ratio** |  | ~1.9 : 1 | ~1.9 : 1 | Calculated |

While these calculations already indicate saturation of carrier protein binding in this dose range, it should be noted that they are necessarily idealized and likely overestimate true binding capacity under biological conditions, where steric constraints, competitive binding, and the heterogeneous composition of CSF further limit effective carrier availability. Thus, it is likely that neither albumin binding nor ApoE-lipoprotein trafficking operate as the dominant distribution mechanism for either conjugate at this dose. Notably, the 10 μg dose used here lies at the lower end of those commonly employed in preclinical mouse studies of CNS ASO delivery. Consequently, saturation of CSF binding partners would be expected to be at least as pronounced, and likely more severe, at the substantially higher doses frequently used. Under these conditions, the majority of the injected dose is therefore predicted to exist as a protein-unbound amphipathic species, which would be expected to associate strongly with lipid-rich surfaces encountered immediately following ICV administration, including the ependymal lining, choroid plexus epithelium, and astrocytic endfeet, through hydrophobicity-driven membrane partitioning. Such avid non-specific membrane associations likely sequester the conjugates at these initial contact surfaces and limit their availability for productive intracellular uptake, consistent with the distribution patterns observed in this study. The biological rationale underpinning lipid conjugation strategies in the systemic compartment therefore does not translate directly to the CSF environment, where the combination of low volume and low protein abundance renders carrier saturation an inherent pharmacokinetic consequence rather than an exceptional condition.

## 4. Conclusion

This study systematically explores a range of lipid-conjugated ASOs, comparing their functional activity and biodistribution across multiple administration routes. When administered subcutaneously, fatty acid conjugates did not redirect ASO activity extrahepatically as intended; rather, they primarily potentiated silencing activity in the liver, with secondary effects observed in the heart and, to a lesser extent, the kidneys, neither of which reached the magnitude of hepatic silencing. The cholesterol conjugate exhibited activity confined to the liver using this route of administration. These results are likely attributable to retention at the injection site, as intravenous administration revealed a broader tissue distribution and higher accumulation of the cholesterol conjugate in the liver and multiple extrahepatic tissues, underscoring the critical role of administration route in determining the pharmacokinetic behavior of lipid-conjugated ASOs.

The majority of ASO studies rely on bulk tissue measurements to infer pharmacodynamic activity, leaving the drug effect at the single-cell level largely unexplored. Here, we applied single-cell sequencing to cells that had taken up lipid-conjugated or unconjugated ASOs, demonstrating that lipid conjugation induces transcriptome-level effects that are not captured by conventional bulk analyses. While the dose used precluded a definitive correlation between cellular uptake and functional target silencing, and therefore did not allow discrimination between productive and non-productive uptake, this work establishes a framework for single-cell resolution analysis of ASO conjugate activity that may inform future mechanistic studies.

To the best of our knowledge, this study is also the first to evaluate activity and biodistribution of lipid-conjugated ASOs following ICV administration. Delivery by this route achieved meaningful silencing in the striatum, though effects in other regions were variable, and potent spinal cord silencing was achieved only with unconjugated ASO. This implies that lipid conjugation does not confer a universal advantage in the CNS and may be detrimental for certain anatomical targets. Biodistribution imaging revealed strikingly increased retention of cholesterol and palmitic acid conjugates relative to unconjugated ASO, consistent with prior reports of rapid ASO clearance from the CNS; however, cholesterol conjugate retention was predominantly localised to membranous structures of the CSF-brain barrier rather than reflecting productive parenchymal distribution. Quantitative estimation of CSF protein binding capacity indicated that the majority of the injected conjugate remains unbound to the principal lipid-carrier proteins at the dose administered, instead associating non-specifically with membrane surfaces surrounding the CSF compartment and potentially limiting productive cellular uptake.

These findings should be interpreted as a proof-of-concept framework established in healthy animals. Pathophysiological conditions relevant to target diseases may naturally alter this distribution landscape through disruption of CSF-brain barrier and BBB integrity, changes in CSF protein composition, and altered systemic lipoprotein and albumin levels, potentially modifying both the carrier saturation constraints estimated here and the access of lipid-conjugated ASOs to parenchymal targets. Validation in disease-relevant models will therefore be essential. Taken together, this study highlights that lipid conjugation strategies must be tailored not only to the molecular target and tissue of interest, but to the physicochemical environment of the delivery compartment itself; a strategy that confers clear benefit in the periphery may be counterproductive in the CNS, and this distinction will need to be carefully considered as lipid-conjugated ASOs advance towards CNS therapeutic applications.

## Abbreviations

ASOs: antisense oligonucleotides
BBB: blood-brain barrier
CNS: central nervous system
CSF: cerebrospinal fluid
DMD: Duchenne muscular dystrophy
FDA: U.S. Food and Drug Administration
FP: fluorescence polarization
HSA: human serum albumin
ICV: intracerebroventricular
IT: intrathecal
IV: intravenous
IVIS: in vivo imaging system
LNA: locked nucleic acid
LSFM: light sheet fluorescence microscopy
NeuN: Neuronal Nuclei
ON: oligonucleotide
PBS: phosphate buffered saline
PMO: phosphorodiamidate morpholino
PS: phosphorothioate
SC: subcutaneous
siRNA: small interfering RNA
TH: Tyrosine Hydroxylase

## Acknowledgments

The authors would like to thank Imaging Specialist Christian Garm and the team at Miltenyi Biotec for uncomplicated technical assistance with everything from 3D sample preparation and the LSFM imaging performed during a demo period of a Ultramicroscope Blaze at Karolinska Institutet. IVIS scans were performed and analyzed at the Preclinical Imaging Facility (PIF) / Preclinical Laboratory (PKL), at Karolinska University Hospital (Sweden), supported by Karolinska Forskning och Utbildning. Histological analysis was performed and analyzed at the P Morphological Phenotype Analysis (FENO), at Karolinska Institutet (Sweden), supported by Karolinska Forskning och Utbildning, Utveckling (FoUU) and Karolinska Institutet infrastructure council. This study was in part performed at the Live Cell Imaging core facility/Nikon Center of Excellence, at Karolinska Institutet, Sweden, supported by the KI infrastructure council and the Olle Engvist Foundation We would also like to acknowledge the contributions of the MedH Flow Cytometry Core Facility financed by the Infrastructure Board at Karolinska Institutet for providing instruments and technical expertise for single cell sorting.

This research was performed under OligoNova initiative funded by Swelife/Vinnova. Further project funding was provided by S.EL.A., supported by the Novo Nordisk Distinguished Innovator Grant, the European Research Council (ERC) under the European Union’s Horizon 2020 Research and Innovation Program (DELIVER, Grant Agreement No. 101001374), the Swedish Foundation of Strategic Research (FormulaEx, SM19-0007), the Swedish Cancer Society (Grant No. 24 3589 Pj 01 H), and the Swedish Research Council (Grant No. 2024-02600) and UK MRC TransNAT. M.H-J was funded by Swedish Research Council (Grant No. 2023-02564).

All figures have been created with Biorender.

## CRediT authorship contribution statement

**Samantha Roudi:** Methodology, Data curation, Investigation, Project Administration, Writing-original draft. **H. Yesid Estupiñán:** Methodology, Data Curation, Investigation, Formal analysis, Visualisation, Writing – review and editing. **Osama Saher:** Methodology, Data curation, Writing – review and editing. **Cristiana Barradas:** Investigation, **Emma Inganäs:** Investigation. **Nicolai Frengen:** Methodology**. Radoslaw Grochowski:** Data curation, Formal Analysis, **Svetlana Pavlova:** Investigation**. Robert Månsson Welinder:** Supervision. **Joel Z. Nordin:** Supervision. **Rula Zain:** Supervision. **Annabelle Biscans:** Methodology, Resources, Writing – review and editing. **Pär Matsson:** Methodology, Data curation, Writing – review and editing. **Rickard Sandberg:** Supervision **Michael Hagemann-Jensen:** Investigation, Formal analysis, Writing – review and editing. **Samir El Andaloussi:** Conceptualization, Funding acquisition, Supervision, Writing-Review and Editing.

## Data Availability

The data that support the findings of this study are available from the corresponding author upon reasonable request.

## References

[1] P. Anand, Y. Zhang, S. Patil, K. Kaur, Metabolic Stability and Targeted Delivery of Oligonucleotides: Advancing RNA Therapeutics Beyond The Liver, J. Med. Chem. 68 (2025) 6870–6896. 10.1021/acs.jmedchem.4c02528.

[2] M. Egli, M. Manoharan, Chemistry, structure and function of approved oligonucleotide therapeutics, Nucleic Acids Res. 51 (2023) 2529–2573. 10.1093/nar/gkad067.

[3] W. Brad Wan, P.P. Seth, The Medicinal Chemistry of Therapeutic Oligonucleotides, J. Med. Chem. 59 (2016) 9645–9667. 10.1021/acs.jmedchem.6b00551.

[4] S.T. Crooke, P.P. Seth, T.A. Vickers, X.H. Liang, The Interaction of Phosphorothioate-Containing RNA Targeted Drugs with Proteins Is a Critical Determinant of the Therapeutic Effects of These Agents, J. Am. Chem. Soc. 142 (2020) 14754–14771. 10.1021/jacs.0c04928.

[5] E. Bäckström, A. Bonetti, P. Johnsson, S. Öhlin, A. Dahlén, P. Andersson, S. Andersson, P. Gennemark, Tissue pharmacokinetics of antisense oligonucleotides, Mol. Ther. Nucleic Acids 35 (2024) 102133. 10.1016/j.omtn.2024.102133.

[6] C. Frank Bennett, H.B. Kordasiewicz, D.W. Cleveland, Antisense Drugs Make Sense for Neurological Diseases, Annu. Rev. Pharmacol. Toxicol. 61 (2021) 831–852. 10.1146/annurev-pharmtox-010919-023738.

[7] C.F. Bennett, A.R. Krainer, D.W. Cleveland, Antisense Oligonucleotide Therapies for Neurodegenerative Diseases, Annual Reviews Neuroscience (2019) 385–406.

[8] B.E. Cook, D.G. Mclaren, J.M. Sullivan, G. El Fakhri, D.L. Yokell, M.W. Freeman, N. Currier, M.E. Oestergaard, H. Dobson, J. Hesterman, N. Salem, I. Nestorov, M. Monine, L. Martarello, K.C. Evans, S. Fradette, T.A. Ferguson, D. Graham, L. Passamonti, Central Nervous System Biodistribution and Pharmacokinetics of Radiolabeled Tofersen in Rodents, Nonhuman Primates, and Humans, J Nucl Med 67 (2026) 139–144. 10.2967/jnumed.125.270731.

[9] FDA, Wainua (eplontersen) FDA package insert, (2023).

[10] FDA, TEGSEDI (inotersen) FDA prescibing information, (2018).

[11] L. Ro, C.R. Calandra, P.A. Kowacs, J.L. Berk, L. Obici, F.A. Barroso, M. Weiler, I. Conceição, S.W. Jung, G. Buchele, M. Brambatti, S.G. Hughes, E. Schneider, N.J. Viney, A. Masri, M.R. Gertz, Y. Ando, J.D. Gillmore, S. Khella, P.J.B. Dyck, M.W. Cruz, N. Investigators, Eplontersen for Hereditary Transthyretin Amyloidosis With Polyneuropathy, JAMA 330 (2023) 1448–1458. 10.1001/jama.2023.18688.

[12] I. Conceição, J.L. Berk, M. Weiler, P.A. Kowacs, N.R. Dasgupta, S. Khella, C.C. Chao, S. Attarian, T.J. Kwoh, S.W. Jung, J. Chen, N.J. Viney, R.Z. Yu, M. Gertz, A. Masri, M.W. Cruz, T. Coelho, Switching from inotersen to eplontersen in patients with hereditary transthyretin-mediated amyloidosis with polyneuropathy: analysis from NEURO-TTRansform, Journal of Neurology 2025 271:10 271 (2024) 6655–6666. 10.1007/s00415-024-12616-6.

[13] C. Ämmälä, W.J. Drury, L. Knerr, I. Ahlstedt, P. Stillemark-Billton, C. Wennberg-Huldt, E.M. Andersson, E. Valeur, R. Jansson-Löfmark, D. Janzén, L. Sundström, J. Meuller, J. Claesson, P. Andersson, C. Johansson, R.G. Lee, T.P. Prakash, P.P. Seth, B.P. Monia, S. Andersson, Targeted delivery of antisense oligonucleotides to pancreatic β-cells, Sci. Adv. 4 (2018) 3386–3403. 10.1126/SCIADV.AAT3386;JOURNAL:JOURNAL:SCIADV;ISSUE:ISSUE:DOI.

[14] S.J. Barker, M.B. Thayer, C. Kim, D. Tatarakis, M.J. Simon, R. Dial, L. Nilewski, R.C. Wells, Y. Zhou, M. Afetian, P. Akkapeddi, A. Chappell, K.S. Chew, J. Chow, A. Clemens, C.B. Discenza, J.C. Dugas, C. Dwyer, T. Earr, C. Ha, Y.S. Ho, D. Huynh, E.I. Lozano, S. Jayaraman, W. Kwan, C. Mahon, M. Pizzo, Y. Robles-Colmenares, E. Roche, L. Sanders, A. Stergioulis, R. Tong, H. Tran, Y.J. Yu Zuchero, A.A. Estrada, K. Gadkar, C.M.M. Koth, P.E. Sanchez, R.G. Thorne, R.J. Watts, T. Sandmann, L.A. Kane, F. Rigo, M.S. Dennis, J.W. Lewcock, S.L. DeVos, Targeting the transferrin receptor to transport antisense oligonucleotides across the mammalian blood-brain barrier, Sci. Transl. Med. 16 (2024) 2245. 10.1126/scitranslmed.adi2245.

[15] O. Sheikh, T. Yokota, Pharmacology and toxicology of eteplirsen and SRP-5051 for DMD exon 51 skipping: an update, Archives of Toxicology 2021 96:1 96 (2021) 1–9. 10.1007/S00204-021-03184-Z.

[16] H. Yin, H.M. Moulton, Y. Seow, C. Boyd, J. Boutilier, P. Iverson, M.J.A. Wood, Cell-penetrating peptide-conjugated antisense oligonucleotides restore systemic muscle and cardiac dystrophin expression and function, Hum. Mol. Genet. 17 (2008) 3909–3918. 10.1093/HMG/DDN293.

[17] M.E. Østergaard, M. Carrer, B.A. Anderson, M. Afetian, M.A. Bakooshli, J.A. Santos, S.K. Klein, J. Capitanio, G.C. Freestone, M. Tanowitz, R. Galindo-Murillo, H.J. Gaus, C.A. Dwyer, M. Jackson, P. Jafar-Nejad, F. Rigo, P.P. Seth, K.U. Gaynor, S.J. Stanway, L. Urbonas, M.A. St. Denis, S. Pellegrino, G.A. Bezerra, M. Rigby, E. Gowans, K. Van Rietschoten, P. Beswick, L. Chen, M.J. Skynner, E.E. Swayze, Conjugation to a transferrin receptor 1-binding Bicycle peptide enhances ASO and siRNA potency in skeletal and cardiac muscles, Nucleic Acids Res. 53 (2025). 10.1093/NAR/GKAF270.

[18] C. Betts, A.F. Saleh, A.A. Arzumanov, S.M. Hammond, C. Godfrey, T. Coursindel, M.J. Gait, M.J. Wood, Pip6-PMO, A New Generation of Peptide-oligonucleotide Conjugates With Improved Cardiac Exon Skipping Activity for DMD Treatment, Mol. Ther. Nucleic Acids 1 (2012) e38. 10.1038/MTNA.2012.30.

[19] Y.Q. Yeoh, A. Amin, B. Cuic, D. Tomas, B.J. Turner, F. Shabanpoor, Efficient systemic CNS delivery of a therapeutic antisense oligonucleotide with a blood-brain barrier-penetrating ApoE-derived peptide, Biomedicine & Pharmacotherapy 175 (2024) 116737. 10.1016/J.BIOPHA.2024.116737.

[20] S.M. Hammond, F. Abendroth, L. Goli, J. Stoodley, M. Burrell, G. Thom, I. Gurrell, N. Ahlskog, M.J. Gait, M.J.A. Wood, C.I. Webster, Antibody-oligonucleotide conjugate achieves CNS delivery in animal models for spinal muscular atrophy, JCI Insight 7 (2022) 1–18. 10.1172/jci.insight.154142.

[21] B. Malecova, R.S. Burke, M. Cochran, M.D. Hood, R. Johns, P.R. Kovach, V.R. Doppalapudi, G. Erdogan, J.D. Arias, B. Darimont, C.D. Miller, H. Huang, A. Geall, H.S. Younis, A.A. Levin, Targeted tissue delivery of RNA therapeutics using antibody– oligonucleotide conjugates (AOCs), Nucleic Acids Res. 51 (2023) 5901–5910. 10.1093/NAR/GKAD415.

[22] S.J. Barker, M.B. Thayer, C. Kim, D. Tatarakis, M.J. Simon, R. Dial, L. Nilewski, R.C. Wells, Y. Zhou, M. Afetian, P. Akkapeddi, A. Chappell, K.S. Chew, J. Chow, A. Clemens, C.B. Discenza, J.C. Dugas, C. Dwyer, T. Earr, C. Ha, Y.S. Ho, D. Huynh, E.I. Lozano, S. Jayaraman, W. Kwan, C. Mahon, M. Pizzo, Y. Robles-Colmenares, E. Roche, L. Sanders, A. Stergioulis, R. Tong, H. Tran, Y.J. Yu Zuchero, A.A. Estrada, K. Gadkar, C.M.M. Koth, P.E. Sanchez, R.G. Thorne, R.J. Watts, T. Sandmann, L.A. Kane, F. Rigo, M.S. Dennis, J.W. Lewcock, S.L. DeVos, Targeting the transferrin receptor to transport antisense oligonucleotides across the mammalian blood-brain barrier, Sci. Transl. Med. 16 (2024) 2245. 10.1126/SCITRANSLMED.ADI2245.

[23] M.E. Østergaard, M. Jackson, A. Low, A. E Chappell, R. G Lee, R.Q. Peralta, J. Yu, G.A. Kinberger, A. Dan, R. Carty, M. Tanowitz, P. Anderson, T.W. Kim, L. Fradkin, A.E. Mullick, S. Murray, F. Rigo, T.P. Prakash, C.F. Bennett, E.E. Swayze, H.J. Gaus, P.P. Seth, Conjugation of hydrophobic moieties enhances potency of antisense oligonucleotides in the muscle of rodents and non-human primates, Nucleic Acids Res. 47 (2019) 6045–6058. 10.1093/nar/gkz360.

[24] T.P. Prakash, A.E. Mullick, R.G. Lee, J. Yu, S.T. Yeh, A. Low, A.E. Chappell, M.E. Østergaard, S. Murray, H.J. Gaus, E. Swayze, P.P. Seth, Fatty acid conjugation enhances potency of antisense oligonucleotides in muscle, Nucleic Acids Res. 47 (2019) 6029–6044. 10.1093/nar/gkz354.

[25] A.E. Chappell, H.J. Gaus, A. Berdeja, R. Gupta, M. Jo, T.P. Prakash, M. Oestergaard, E.E. Swayze, P.P. Seth, Mechanisms of palmitic acid-conjugated antisense oligonucleotide distribution in mice, Nucleic Acids Res. 48 (2020) 4382–4395. 10.1093/NAR/GKAA164.

[26] S. Ait Benichou, D. Jauvin, T. De Serres-Bérard, F. Bennett, F. Rigo, G. Gourdon, M. Boutjdir, M. Chahine, J. Puymirat, Enhanced Delivery of Ligand-Conjugated Antisense Oligonucleotides ( C16-HA-ASO ) Targeting Dystrophia Myotonica Protein Kinase Transcripts for the Treatment of Myotonic Dystrophy Type 1, https://Home.Liebertpub.Com/Hum 33 (2022) 810–820. 10.1089/hum.2022.069.

[27] A. Biscans, A. Coles, R. Haraszti, Di. Echeverria, M. Hassler, M. Osborn, A. Khvorova, Diverse lipid conjugates for functional extra-hepatic siRNA delivery in vivo, Nucleic Acids Res. 47 (2019) 1082–1096. 10.1093/nar/gky1239.

[28] Ö. Kartal, F. Andres, M.P. Lai, R. Nehme, K. Cottier, waveRAPID—A Robust Assay for High-Throughput Kinetic Screens with the Creoptix WAVEsystem, SLAS Discovery 26 (2021) 995–1003. 10.1177/24725552211013827.

[29] J.M. Chamouard, J. Barre, S. Urien, G. Houin, J.P. Tillement, Diclofenac binding to albumin and lipoproteins in human serum, Biochem. Pharmacol. 34 (1985) 1695–1700. 10.1016/0006-2952(85)90636-7.

[30] Ö. Kartal, F. Andres, M.P. Lai, R. Nehme, K. Cottier, waveRAPID—A Robust Assay for High-Throughput Kinetic Screens with the Creoptix WAVEsystem, SLAS Discovery 26 (2021) 995–1003. 10.1177/24725552211013827.

[31] K. Rhéaume, Z. Chen, Y. Wang, C. Plante, D.H. Bostanthirige, M. Lévesque, S. Geha, L.Q. Le, J.P. Brosseau, Whole Central and Peripheral Nervous System Mice Dissection, J. Vis. Exp. 2023 (2023). 10.3791/64974.

[32] Miltenyi Biotec, Immunostaining and clearing of mouse brain hemispheres with preservation of endogenous fluorescent protein signal, (n.d.). https://www.miltenyibiotec.com/SE-en/applications/all-protocols/immunostaining-and-clearing-of-mouse-brain-hemispheres-with-preservation-of-endogenous-fluorescent-protein-signal.html?query=:relevance:codeString:130-090-753:codeString:130-126-335:codeStrin.

[33] M. Hagemann-Jensen, C. Ziegenhain, P. Chen, D. Ramsköld, G.J. Hendriks, A.J.M. Larsson, O.R. Faridani, R. Sandberg, Single-cell RNA counting at allele and isoform resolution using Smart-seq3, Nature Biotechnology 2020 38:6 38 (2020) 708–714. 10.1038/s41587-020-0497-0.

[34] M.E. Østergaard, M. Jackson, A. Low, A.E. Chappell, R.G. Lee, R.Q. Peralta, J. Yu, G.A. Kinberger, A. Dan, R. Carty, M. Tanowitz, P. Anderson, T. Kim, L. Fradkin, A.E. Mullick, S. Murray, F. Rigo, T.P. Prakash, C.F. Bennett, E.E. Swayze, H.J. Gaus, P.P. Seth, Conjugation of hydrophobic moieties enhances potency of antisense oligonucleotides in the muscle of rodents and non-human primates, Nucleic Acids Res. 47 (2019) 6045–6058. 10.1093/nar/gkz360.

[35] E.A. Kusznir, J.C. Hau, M. Portmann, A.G. Reinhart, F. Falivene, J. Bastien, J. Worm, A. Ross, M. Lauer, P. Ringler, F. Sladojevich, S. Huber, K. Bleicher, M. Keller, Propensities of Fatty Acid-Modified ASOs: Self-Assembly vs Albumin Binding, Bioconjug. Chem. 34 (2023) 866–879. 10.1021/acs.bioconjchem.3c00085.

[36] A. Khvorova, M. Nikan, M. Hassler, M. Osborn, R. Haraszti, A. Coles, A. Turanov, N. Aronin, Bioactive conjugates for oligonucleotide delivery, US10633653B2, 2016.

[37] R.Z. Yu, R. Gunawan, Z. Li, R.S. Mittleman, A. Mahmood, J.S. Grundy, W. Singleton, No effect on QT intervals of mipomersen , a 2 ′ - O -methoxyethyl modified antisense oligonucleotide targeting ApoB-100 mRNA , in a phase I dose escalation placebo-controlled study , and confirmed by a thorough QT ( tQT ) study , in healthy subjects, (2016) 267–275. 10.1007/s00228-015-1992-y.

[38] A. Biscans, A. Coles, R. Haraszti, Di. Echeverria, M. Hassler, M. Osborn, A. Khvorova, Diverse lipid conjugates for functional extra-hepatic siRNA delivery in vivo, Nucleic Acids Res. (2019). 10.1093/nar/gky1239.

[39] M.F. Osborn, A.H. Coles, A. Biscans, R.A. Haraszti, L. Roux, S. Davis, S. Ly, D. Echeverria, M.R. Hassler, B.M.D.C. Godinho, M. Nikan, A. Khvorova, Hydrophobicity drives the systemic distribution of lipid-conjugated siRNAs via lipid transport pathways, 47 (2019) 1070–1081. 10.1093/nar/gky1232.

[40] S. Wada, H. Yasuhara, F. Wada, M. Sawamura, R. Waki, Evaluation of the effects of chemically different linkers on hepatic accumulations, cell tropism and gene silencing ability of cholesterol-conjugated antisense oligonucleotides, Journal of Controlled Release 226 (2016) 57–65. 10.1016/j.jconrel.2016.02.007.

[41] P. Mukherjee, E. Aksamitiene, A. Alex, J. Shi, K. Bera, C. Zhang, D.R. Spillman, M. Marjanovic, M. Fazio, P.P. Seth, K. Frazier, S.R. Hood, S.A. Boppart, Differential Uptake of Antisense Oligonucleotides in Mouse Hepatocytes and Macrophages Revealed by Simultaneous Two-Photon Excited Fluorescence and Coherent Raman Imaging, Nucleic Acid Ther. 32 (2022) 163–176. 10.1089/NAT.2021.0059;WEBSITE:WEBSITE:SAGE;ISSUE:ISSUE:DOI

[42] B. Spencer-Dene, P. Mukherjee, A. Alex, K. Bera, W.J. Tseng, J. Shi, E.J. Chaney, D.R. Spillman, M. Marjanovic, E. Miranda, S.A. Boppart, S.R. Hood, Localization of unlabeled bepirovirsen antisense oligonucleotide in murine tissues using in situ hybridization and CARS imaging, RNA 29 (2023) 1575–1590. 10.1261/RNA.079699.123/-/DC1.

[43] M. Tanowitz, L. Hettrick, A. Revenko, G.A. Kinberger, T.P. Prakash, P.P. Seth, Asialoglycoprotein receptor 1 mediates productive uptake of N-acetylgalactosamine-conjugated and unconjugated phosphorothioate antisense oligonucleotides into liver hepatocytes, Nucleic Acids Res. 45 (2017) 12388–12400. 10.1093/NAR/GKX960.

[44] H. Yasuhara, K. Kadotsuji, K. Watanabe, T. Kakutani, T. Tochitani, I. Mise, M. Konishi, T. Nakagawa, I. Miyawaki, Unveiling Liver Micro-Distribution: NanoSIMS Imaging Reveals Critical Intracellular Distribution of Chemically Modified Antisense Oligonucleotides, Nucleic Acid Ther. (2025). 10.1177/21593337251399181;SUBPAGE:STRING:ACCESS.

[45] T.C. Roberts, M.J.A. Wood, K.E. Davies, Therapeutic approaches for Duchenne muscular dystrophy, Nature Reviews Drug Discovery 2023 22:11 22 (2023) 917–934. 10.1038/s41573-023-00775-6.

[46] T. Metz, M.M. Welling, E. Suidgeest, E. Nieuwenhuize, T. de Vlaam, D. Curtis, T.T. Hailu, L. van der Weerd, W.M.C. van Roon-Mom, Biodistribution of Radioactively Labeled Splice Modulating Antisense Oligonucleotides After Intracerebroventricular and Intrathecal Injection in Mice, Nucleic Acid Ther. 34 (2024) 26–34. 10.1089/nat.2023.0018.

[47] P. Jafar-Nejad, B. Powers, A. Soriano, H. Zhao, D.A. Norris, J. Matson, B. Debrosse-Serra, J. Watson, P. Narayanan, S.J. Chun, C. Mazur, H. Kordasiewicz, E.E. Swayze, F. Rigo, The atlas of RNase H antisense oligonucleotide distribution and activity in the CNS of rodents and non-human primates following central administration, Nucleic Acids Res. 49 (2021) 657–673. 10.1093/nar/gkaa1235.

[48] A. Korff, X. Yang, O. Ozdemir, A. Samanta, Y.-D. Wang, T. Patni, A.J. Lavado, A.M. Kavirayani, J. Ochaba, B. Powers, C.F. Bennett, H.J. Kim, J.P. Taylor, Preclinical evaluation of antisense oligonucleotide therapy in a mouse model of HNRNPH2-related neurodevelopmental disorder, BioRxiv (2025) 2025.11.04.686541. 10.1101/2025.11.04.686541.

[49] A.G. Sorets, K.R. Schwensen, N. Francini, A. Kjar, A.M. Abdulrahman, A. Shostak, K.A. Katdare, K.M. Schoch, R.P. Cowell, J.C. Park, A.P. Ligocki, W.T. Ford, L. Ventura-Antunes, E.N. Hoogenboezem, A. Prusky, M. Castleberry, D.L. Michell, E. Fritsch, S.M. Lyons, T.M. Miller, K.C. Vickers, M.S. Schrag, C.L. Duvall, E.S. Lippmann, Elucidating brain transport pathways and cell type-dependent gene silencing of a durable lipid–siRNA conjugate administered into cerebrospinal fluid, Nucleic Acids Res. 53 (2025). 10.1093/nar/gkaf600.

[50] C. Mazur, B. Powers, K. Zasadny, J.M. Sullivan, H. Dimant, F. Kamme, J. Hesterman, J. Matson, M. Oestergaard, M. Seaman, R.W. Holt, M. Qutaish, I. Polyak, R. Coelho, V. Gottumukkala, C.M. Gaut, M. Berridge, N.J. Albargothy, L. Kelly, R.O. Carare, J. Hoppin, H. Kordasiewicz, E.E. Swayze, A. Verma, Brain pharmacology of intrathecal antisense oligonucleotides revealed through multimodal imaging, JCI Insight 4 (2019) e129240. 10.1172/JCI.INSIGHT.129240.

[51] M. Monine, D. Norris, Y. Wang, I. Nestorov, A physiologically-based pharmacokinetic model to describe antisense oligonucleotide distribution after intrathecal administration, J. Pharmacokinet. Pharmacodyn. 48 (2021) 639–654. 10.1007/s10928-021-09761-0.

[52] M. Butler, C.S. Hayes, A. Chappell, S.F. Murray, T.L. Yaksh, X.Y. Hua, Spinal distribution and metabolism of 2′-O-(2-methoxyethyl)-modified oligonucleotides after intrathecal administration in rats, Neuroscience 131 (2005) 705–715. 10.1016/J.NEUROSCIENCE.2004.11.038.

[53] A. Mohan, B. Fitzsimmons, H.T. Zhao, Y. Jiang, C. Mazur, E.E. Swayze, H.B. Kordasiewicz, Antisense oligonucleotides selectively suppress target RNA in nociceptive neurons of the pain system and can ameliorate mechanical pain, Pain 159 (2018) 139–149. 10.1097/J.PAIN.0000000000001074.

[54] T. Berkman, X. Li, Y. Liang, A. Korban, A. Bekker, Y.X. Tao, Systemic administration of NIS-lncRNA antisense oligonucleotide alleviates neuropathic pain, Neurosci. Lett. 817 (2023) 137512. 10.1016/J.NEULET.2023.137512.

[55] C.H. Wen, T. Berkman, X. Li, S. Du, G. Govindarajalu, H. Zhang, A. Bekker, S. Davidson, Y.X. Tao, Effect of intrathecal NIS-lncRNA antisense oligonucleotides on neuropathic pain caused by nerve trauma, chemotherapy, or diabetes mellitus, Br. J. Anaesth. 130 (2023) 202–216. 10.1016/J.BJA.2022.09.027.

[56] S. Whish, K.M. Dziegielewska, K. Møllgård, N.M. Noor, S.A. Liddelow, M.D. Habgood, S.J. Richardson, N.R. Saunders, The inner csf-brain barrier: Developmentally controlled access to the brain via intercellular junctions, Front. Neurosci. 9 (2015) 124778. 10.3389/FNINS.2015.00016/TEXT.

[57] W.M. Pardridge, CSF, blood-brain barrier, and brain drug delivery, Expert Opin. Drug Deliv. 13 (2016) 963–975. 10.1517/17425247.2016.1171315.

[58] S. Curry, H. Mandelkow, P. Brick, N. Franks, Crystal structure of human serum albumin complexed with fatty acid reveals an asymmetric distribution of binding sites, Nature Structural Biology 1998 5:9 5 (1998) 827–835. 10.1038/1869.

[59] I. Petitpas, T. Grüne, A.A. Bhattacharya, S. Curry, Crystal structures of human serum albumin complexed with monounsaturated and polyunsaturated fatty acids, J. Mol. Biol. 314 (2001) 955–960. 10.1006/JMBI.2000.5208.

[60] J.D. Ulrich, J.M. Burchett, J.L. Restivo, D.R. Schuler, P.B. Verghese, T.E. Mahan, G.E. Landreth, J.M. Castellano, H. Jiang, J.R. Cirrito, D.M. Holtzman, In vivo measurement of apolipoprotein E from the brain interstitial fluid using microdialysis, Mol. Neurodegener. 8 (2013) 13. 10.1186/1750-1326-8-13.

[61] J.P. Segrest, M.C. Cheung, M.K. Jones, Volumetric determination of apolipoprotein stoichiometry of circulating HDL subspecies, J. Lipid Res. 54 (2013) 2733. 10.1194/JLR.M039172.

[62] J.B. Massey, H.J. Pownall, S. Macha, J. Morris, M.R. Tubb, R.A.G.D. Silva, Mass spectrometric determination of apolipoprotein molecular stoichiometry in reconstituted high density lipoprotein particles, J. Lipid Res. 50 (2009) 1229. 10.1194/JLR.D800044-JLR200.

[63] Y. Qi, J. Fan, J. Liu, W. Wang, M. Wang, J. Sun, J. Liu, W. Xie, F. Zhao, Y. Li, D. Zhao, Cholesterol-overloaded HDL particles are independently associated with progression of carotid atherosclerosis in a cardiovascular disease-free population: A community-based cohort study, J. Am. Coll. Cardiol. 65 (2015) 355–363. 10.1016/j.jacc.2014.11.019.

[64] A.T. Remaley, HDL cholesterol/HDL particle ratio: A new measure of HDL function?, J. Am. Coll. Cardiol. 65 (2015) 364–366. 10.1016/j.jacc.2014.11.018.

[65] A. Kontush, M. Lhomme, M.J. Chapman, Unraveling the complexities of the HDL lipidome, J. Lipid Res. 54 (2013) 2950–2963. 10.1194/jlr.r036095.

