## Supplementary Data for "Lipid–ASO therapeutics exhibit differential tissue targeted delivery upon systemic or local CNS administration"

### Supplementary information

| No | Name | Sequence 5'-3' | Abs. Coeff. (L/mol/cm) | MW (g/mol) |
|---|---|---|---|---|
| 1 | EPA-ASO | EPA-HA***G*mC*A***T*T*mC*T*A*A*T*A*G*mC***A*G*mC** | 152800 | 5809.97 |
| 2 | Palm-ASO | Palm-HA***G*mC*A***T*T*mC*T*A*A*T*A*G*mC***A*G*mC** | 152800 | 5763.94 |
| 3 | DCA-ASO | DCA-HA***G*mC*A***T*T*mC*T*A*A*T*A*G*mC***A*G*mC** | 152800 | 5848.10 |
| 4 | Chol-ASO | Chol-HA***G*mC*A***T*T*mC*T*A*A*T*A*G*mC***A*G*mC** | 152800 | 5938.18 |
| 5 | Chol- scramble ASO | Chol-HA***G*G*mC***mC*A*A*T*A*mC*G*mC*mC*G***T*mC*A** | 146100 | 5952.21 |
| 6 | ASO | **G*mC*A***T*T*mC*T*A*A*T*A*G*mC***A*G*mC** | 152800 | 5326.55 |
| 7 | ASO-Cy3 | **G*mC*A***T*T*mC*T*A*A*T*A*G*mC***A*G*mC***Cy3 | 149400 | 5853.96 |
| 8 | Chol-ASO-Cy3 | Chol-HA***G*mC*A***T*T*mC*T*A*A*T*A*G*mC***A*G*mC***Cy3 | 149400 | 6461.83 |
| 9 | Palm-ASO-Cy3 | Palm-HA***G*mC*A***T*T*mC*T*A*A*T*A*G*mC***A*G*mC***Cy3 | 149400 | 6287.59 |
| 10 | HA-ASO | HA* **G*mC*A***T*T*mC*T*A*A*T*A*G*mC***A*G*mC** | 152800 | 5525.53 |

**Supplementary Table 1: Analytical data for lipid-ASO conjugates.** The backbone of ASOs is fully PS-modified, and the ASO design is 3-10-3, where ten nucleotides in the middle are DNA, spanned by three locked nucleic acids (LNAs) on each end (in bold blue). mC denotes modification of cytosine base to 5-methylcytosine. HA=hexylamine, EPA=Eicosapentaenoic Acid, Palm=Palmitic Acid, DCA= Docosanoic Acid, Chol=Cholesterol

| No | Name | KD (side 1) µM | KD (side 2) µM | MTR (side 1) s | MTR (side 2) s |
|---|---|---|---|---|---|
| 1 | EPA-ASO | 12.64 (± 3.98) | 13.10 (± 5.24) | 0.23 (± 3.99) | 0.16 (± 0.13) |
| 2 | Palm-ASO | 2.81 (± 1.35) | 6.87 (± 1.83) | 1.55 (± 1.52) | 0.29 (± 0.06) |
| 3 | DCA-ASO | 0.52 (± 1.56) | 1.96 (± 1.42) | 18.15 (±1.42) | 1.49 (±0.25) |
| 4 | Chol-ASO | 1.49 (± 3.34) | 6.55 (± 2.65) | 28.36 (± 3.96) | 0.89 (± 0.37) |
| 5 | Chol-scramble ASO | 0.69 (± 5.85) | 6.12 (± 3.21) | 26.45 (± 5.47) | 1.01 (± 0.38) |
| 6 | ASO | No MB | | No MB | |
| 7 | ASO-Cy3 | No MB | | No MB | |
| 8 | Chol-ASO-Cy3 | 2.72 (± 2.54) | 4.11 (± 2.05) | 18.91 (± 1.37) | 1.54 (± 0.13) |
| 9 | Palm-ASO-Cy3 | 1.54 (± 1.13) | 2.47 (± 1.22) | 3.65 (± 1.38) | 0.39 (± 0.11) |

No MB: No measurable binding

**Supplementary Table 2. Calculated dissociation constant (Kd) and mean residence time (MRT) for lipid-ASOs binding to HSA.** Two-site binding model was applied, e.g. high-affinity and low-affinity based on diclofenac reference. KD values inform the affinity of the molecule, since it depends on binding and dissociation rate ($K_D = k_{off} / k_{on}$), whereas the MRT is a function of dissociation rate only (MRT= $1/k_{off}$).

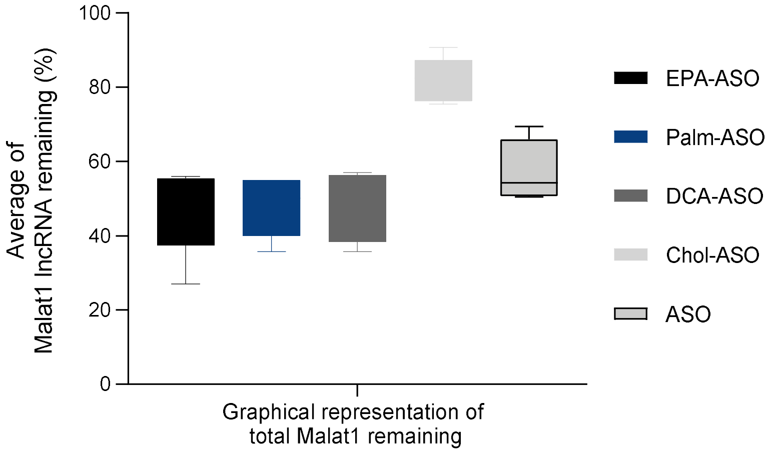


**Supplementary Figure 1. Overall activity of lipid-ASOs or ASO upon subcutaneous administration.** Graphical representation of cumulative activity of each conjugate, calculated by averaging the percentage of Malat1 lnRNA remaining in all organs determined upon subcutaneous administration of lipid-ASOs or ASO only.

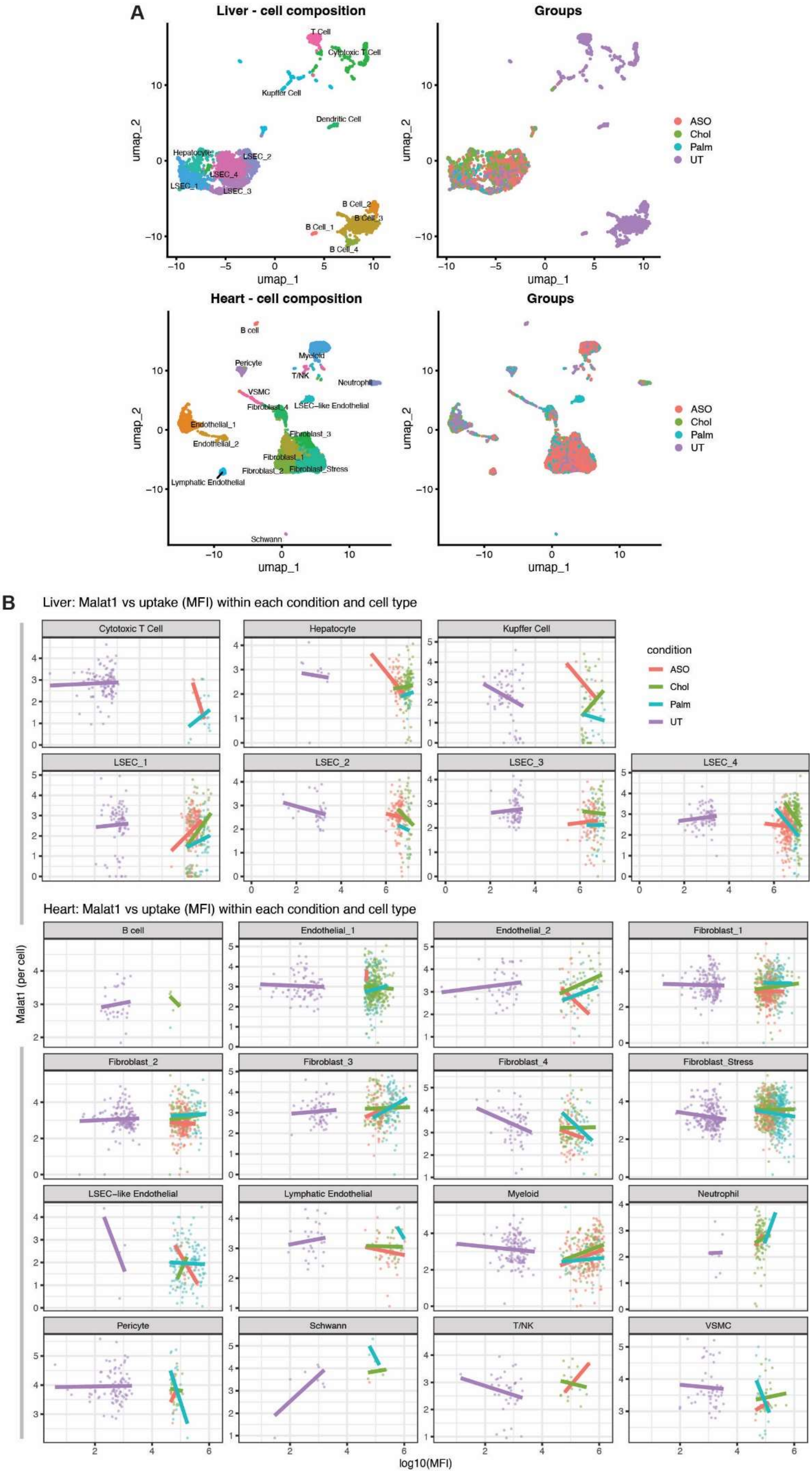
A
Liver - cell composition
Groups
umap_1
umap_2
T Cell
Cytotoxic T Cell
Kupffer Cell
Dendritic Cell
Hepatocyte
LSEC_1
LSEC_2
LSEC_3
LSEC_4
B Cell_1
B Cell_2
B Cell_3
B Cell_4
ASO
Chol
Palm
UT
Heart - cell composition
B cell
Pericyte
Myeloid
T/NK
Neutrophil
VSMC
Fibroblast_4
LSEC–like Endothelial
Endothelial_1
Endothelial_2
Fibroblast_3
Fibroblast_1
Fibroblast_2
Fibroblast_Stress
Lymphatic Endothelial
Schwann
B
Liver: Malat1 vs uptake (MFI) within each condition and cell type
condition
Heart: Malat1 vs uptake (MFI) within each condition and cell type
Malat1 (per cell)
log10(MFI)

**Supplementary Figure 2. Single-cell clustering and assessment of productive versus non-productive uptake of Cy3-labeled ASOs. A)** Uniform Manifold Approximation and Projection (UMAP) visualization of single cells isolated from the liver (top) and heart (bottom), divided by cell type (left) or colored by treatment group (right). **B)** Assessment of productive versus non-productive uptake at the single-cell level by integrating *Malat1* transcript abundance with the corresponding Cy3 mean fluorescence intensity (MFI) recorded during fluorescence-activated cell sorting. Data are shown for liver (top) and heart (bottom) samples.

**A)** Video SC ASO-Cy3.mp4 **B)** Video SC Chol-ASO-Cy3.mp4 **C)** Video SC Palm-ASO-Cy3.mp4

**Supplementary Video 1. Three-dimensional biodistribution of Cy3-labeled ASOs in the spinal cord following intracerebroventricular administration.** Z-stack reconstruction of the spinal cord 48 h after ICV administration of 10 µg Cy3-labeled ASO. Cy3 signal is shown in green, while NeuN and tyrosine hydroxylase (TH) immunostaining are shown in red. **A)** ASO-Cy3, **B)** Chol-ASO-Cy3, and **C)** Palm-ASO-Cy3.

**A)** Video lumbar DRG ASO-Cy3.mp4 **B)** Video lumbar SC Chol-ASO-Cy3.mp4

**Supplementary Video 2. Three-dimensional biodistribution of Cy3-labeled ASOs in the lumbar spinal cord following intracerebroventricular administration.** Three-dimensional reconstruction of the lumbar spinal cord 48 h after intracerebroventricular (ICV) administration of 10 µg Cy3-labeled ASO, highlighting dorsal root ganglia (DRG) and their co-localization with neuronal markers. Cy3 fluorescence is shown in green, while NeuN and tyrosine hydroxylase (TH) immunostaining are shown in red. **A)** ASO-Cy3 and **B)** Chol-ASO-Cy3.
